# CDKN1A is a target for phagocytosis-mediated cellular immunotherapy in acute leukemia

**DOI:** 10.1101/2022.01.18.476736

**Authors:** Awatef Allouch, Laurent Voisin, Yanyan Zhang, Raza Syed, Yann Lecluse, Julien Calvo, Dorothée Sélimoglu-Buet, Stéphane De Botton, Fawzia Louache, Françoise Pflumio, Eric Solary, Jean-Luc Perfettini

## Abstract

Most tumor-associated macrophages (TAMs), which are abundant in the tumor microenvironment, demonstrate an immunosuppressive phenotype and contribute to tumor progression, treatment resistance and poor clinical outcomes^1,2^. Due to their functional plasticity, these cells could be reprogrammed to acquire a pro-inflammatory phenotype and promote tumor clearance^1^. Several therapeutic approaches targeting TAMs to alleviate their immunosuppressive properties or to harness their tumoricidal capacities have been developed^3,4^. Inhibition of interactions between phagocytic inhibitor receptors on macrophages and “don’t eat me signals” on cancer cells, which promotes cancer cell engulfment, showed therapeutic benefits for several tumor types^4–6^. Investigating mechanisms involved in macrophage-mediated phagocytosis of tumor cells, we demonstrate here a key role for the cyclin-dependent kinase inhibitor CDKN1A (p21). Through transcriptional repression of *SIRPα* (Signal-Regularity Protein α), which encodes a phagocytic inhibitor, CDKN1A promotes the ability of monocyte-derived macrophages (MDMs) to engulf leukemic cells. In turn, these MDMs acquire a pro-inflammatory phenotype that extends to surrounding MDMs in an Interferon γ (IFNγ)-dependent manner. Human monocytes genetically engineered to overexpress p21 (p21TD-Mo) differentiate into anti-inflammatory MDMs that are primed for leukemic cell phagocytosis when transferred into mice xenografted with patient-derived T-cell acute lymphoblastic leukemia (T-ALL) cells. After leukemic cell engulfment, engineered macrophages undergo a pro-inflammatory activation, reducing leukemic burden and substantially prolonging survival of mice. These results reveal p21 as a trigger of phagocytosis-guided pro-inflammatory reprogramming of TAMs and demonstrate the potential for p21TD-Mo-based cell therapy in cancer immunotherapy.

## Main text

Phagocytosis of cancer cells by macrophages plays a critical role in cancer immunosurveillance^7,8^. Cancer cells can evade macrophage-mediated phagocytosis by up-regulating “don’t eat me signals” on their surfaces such as CD47, PDL-1, β2M and CD24, which bind to the phagocytic inhibitor receptors SIRPα, PD-1, LILRB1 and Siglec-10, respectively^7,9–12^. These interactions trigger intracellular cascades of inhibitory signals in macrophages to block cytoskeletal rearrangements, the formation of phagocytic synapses and the engulfment of cancer cells^7,9–13^. Several immunotherapies aim to disrupt these interactions, through the use of macrophage immune checkpoint blockers (MICB) (such as blocking antibodies or antagonist engineered SIRPα variants), to circumvent negative signaling of phagocytosis, enabling macrophages to engulf and clear cancer cells^7,9–12,14,15^. However, the resistance of various cancer cells to the MICB reveals the existence of yet unknown regulatory mechanisms of tumor phagocytosis.

Human blood MDMs, which possess an anti-inflammatory phenotype^16^, were co-cultured with several leukemic cell lines (Fig. 1a). MDMs rapidly engulfed Jurkat T cells (Fig. 1b) and delivered them to lysosomal compartments for degradation, as visualized by confocal microscopy (Extended Data Fig. 1a). MDMs demonstrated phagocytic activity toward two other T-ALL cell lines (MOLT4 or CEM) without engulfment of primary peripheral blood lymphocytes (PBLs) (Fig. 1c). In these co-culture conditions, MDMs also engulfed acute myeloid leukemia (AML) cells such as the THP1 cell line (Fig. 1c) and primary AML blast cells (Fig. 1d) (Extended Data Table1), but failed to engulf erythroleukemic HEL cells and chronic myelogenous leukemic K562 cells (Fig. 1c), indicating that MDMs engulfed preferentially acute leukemia cells. In accordance with the previously demonstrated role of ROCK-dependent signaling pathways in phagocytosis^17^, MDM engulfment of leukemic cells and cell lines may depend on ROCK activity, as the pharmacological inhibition of ROCK kinase activity with Y27632 (as revealed by the inhibition of myosin light chain 2 phosphorylation on serine 19 (MLC2S19*) (Extended Data Fig. 1b)) abolished Jurkat cell phagocytosis (Extended Data Fig. 1c). In contrast, neither the broad spectrum-caspase inhibitor (ZVAD) (Fig. 1c,d) nor recombinant human annexin V (Extended Data Fig. 1d,e), which respectively impair apoptosis and the uptake of dying cells by macrophages, reduced the engulfment of Jurkat cells and primary AML blast cells, thus excluding efferocytosis^18^. We also excluded a role for target cell geometry since no correlation was identified between the phagocytosis of leukemic cells and the target cell’s volume and surface area as determined by confocal microscopy (Extended Data Fig. 1f,g).

**Fig. 1.**
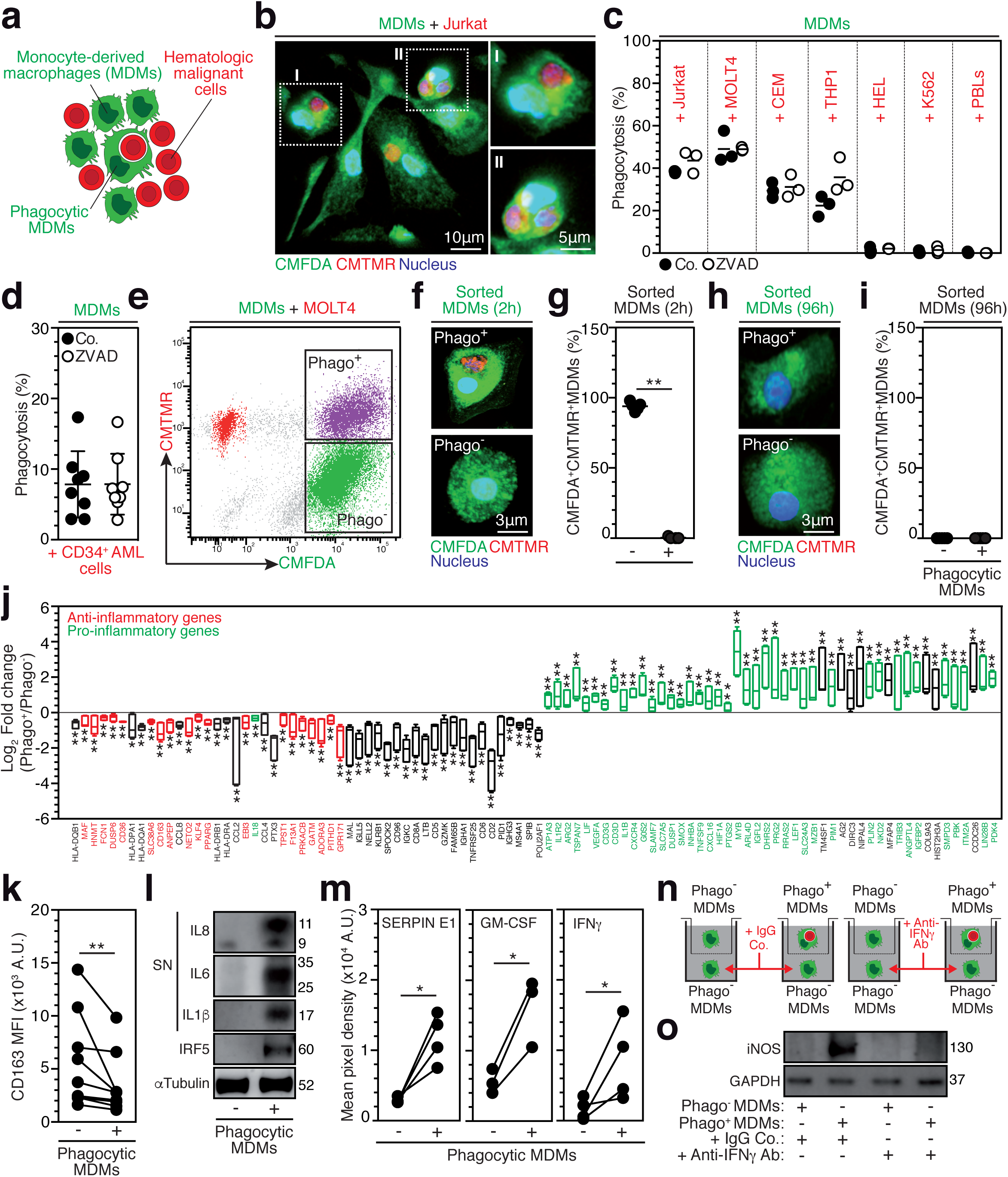
Tumor phagocytosis triggers the pro-inflammatory activation of macrophages. **a**, Co-culture model of CMFDA-labeled MDMs and CMTMR-labeled hematologic malignant cells. **b,** Confocal micrograph of MDMs and Jurkat cells after 8 hours (8h) of co-culture. **c**,**d**, Percentage of phagocytosis, determined by the number of MDMs engulfing leukemic cells or PBLs (**c**) or AML CD34^+^blasts (**d**) divided on total MDMs (at least 100 MDMs in five fields) in control (Co.) or ZVAD (100 µM) treated co-cultures. **e**, FACS dot plot of Phago^+^MDMs or Phago^-^MDMs sorting after 2h of co-culture with MOLT4 cells. **f-g**, Confocal micrographs and percentages of CMFDA^+^CMTMR^+^MDMs 2h (**p=0.0079) (**f**,**g**) and 96h (**h**,**i**) after the sorting of Phago^+^MDMs and Phago^-^MDMs. **j-m,** Phago^+^MDMs were analyzed, as compared to phago^-^MDMs, for modulated genes by microarray (**p=0.0022, *p=0.0152) (**j**), CD163 membrane expression by FACS (**p=0.0039) (**k**), IRF5 expression by western blot (WB) (**l**) and supernatant (SN) indicated pro-inflammatory cytokines by WB (**l**) or by cytokine microarray (*p=0.0143, *p=0.05, *p=0.0286, respectively) (**m**) at 96h (**j**,**k**,**m**) and 7d (**l**) after FACS sorting. **n,o,** Transwell co-cultures model of Phago^+^MDMs and Phago^-^MDMs at 2h after FACS sorting (**n**) and iNOS expression by WB of Phago^-^MDMs co-cultured in the bottom chambers during 15d (**o**). In (**b,l,o**) and **(e,f,h**) data are representative of n=3 and n=5 donors. In (**c,j**), (**d)** and (**g,i**) data are means±SEM from n=3, n=6 and n=5 donors. In (**d)** CD34^+^ cells are from n=4 AML patients. In (**k)** and (**m)** data are donor-matched from n=9 and in (**m)** n=4 (for SERPIN E1, IFNγ) and n=3 (for GM-CSF). *p<0.05, **p<0.01 are determined with two- (**g)** or one- (**m)** tailed unpaired Mann-Whitney, Kolmogorov-Smirnov (**j)** and two-tailed paired Wilcoxon (**k),** tests.

In order to explore the functional impact of leukemic cell phagocytosis on MDM phenotype, we cultured MDMs with MOLT4 cells for 2 hours before sorting phagocytic (Phago^+^MDMs) and non-phagocytic (Phago^-^MDMs) cells. Phago^+^MDMs (Fig. 1e-g) showed a complete degradation of engulfed cells 96 hours after cell sorting (Fig. 1h,i). Gene expression analysis indicated that, after degradation of target cells, Phago^+^MDMs underwent pro-inflammatory activation (Fig. 1j and Extended Data Table 2), as indicated by the up-regulation of 40 pro-inflammatory genes and down-regulation of 19 anti-inflammatory genes, compared with Phago^-^MDMs (Fig. 1j, Extended Data Table 2). Consistently, Phago^+^MDMs demonstrated decreased expression of the cell surface scavenger receptor CD163 (Fig. 1k), increased expression of the transcription factor IRF5 (Interferon regulatory factor 5, Fig. 1l), and enhanced release of pro-inflammatory cytokines including IL1β, IL6, IL8, SERPIN E1, GM-CSF, IFNγ, IL23, Gro-α, IL1-ra, MIF and IL27 (Fig. 1l,m and Extended Data Fig. 2a), compared with Phago^-^MDMs. Similarly, Phago^+^MDMs that have engulfed primary AML blast cells demonstrated increased secretion of IL8, compared with Phago^-^MDMs (Extended Data Fig. 2b).

Considering that Interferon γ (IFNγ), which was secreted by Phago^+^MDMs (Fig. 1m), can potentially modulate the expression of 53 target genes in Phago^+^MDMs (Extended Data Table 2) and convert immunosuppressive TAMs into immunostimulatory macrophages^19,20^, we next explored the ability of secreted IFNγ to promote the proinflammatory reprogramming of surrounding Phago^-^MDMs. Phago^+^MDMs in the upper chamber of trans-well devices were co-cultured with Phago^-^MDMs in the bottom chamber, in the absence or presence of control IgG or anti-IFNγ blocking antibodies for 15 days (Fig. 1n). While increased expression of the pro-inflammatory inducible oxide synthase (iNOS) was detected in Phago^-^MDMs in the presence of control IgG, this effect was lost in the presence of anti-INFγ-blocking antibodies (Fig. 1o), demonstrating that Phago^+^MDMs could subsequently convert bystander anti-inflammatory MDMs to a pro-inflammatory phenotype.

In order to identify molecular mechanisms regulating the macrophage-mediated phagocytosis of leukemic cells, we assessed the role of p21, which has been involved in differentiation and survival of macrophages^21–24^. Using specific small interfering RNA, MDMs were depleted for p21 (Fig. 2a) and assessed for phagocytosis. Interestingly, p21 depletion strongly reduced phagocytosis of Jurkat cells (Fig. 2b,c), MOLT4 cells (Fig. 2d) and primary human AML blast cells (Fig. 2e), without affecting the phagocytosis of pHrodo green bacterial *E. coli* bioparticles (Fig. 2f). These results suggested that p21 is a key regulator of the MDM-mediated phagocytosis of leukemic cells. Inversely, increased p21 expression (Extended Data Fig. 3a,b,f) by treatment of MDMs with phorbol myristate acetate (PMA) ^25^, histone deacetylase (HDAC) inhibitor MS275 ^26^ or intravenous immunoglobulins (IVIg) ^16^, significantly enhanced the phagocytosis of Jurkat cells and MOLT4 cells (Extended Data Fig. 3c-e,g), confirming that p21 is a master regulator of MDM-mediated phagocytosis of leukemic cells.

**Fig. 2.**
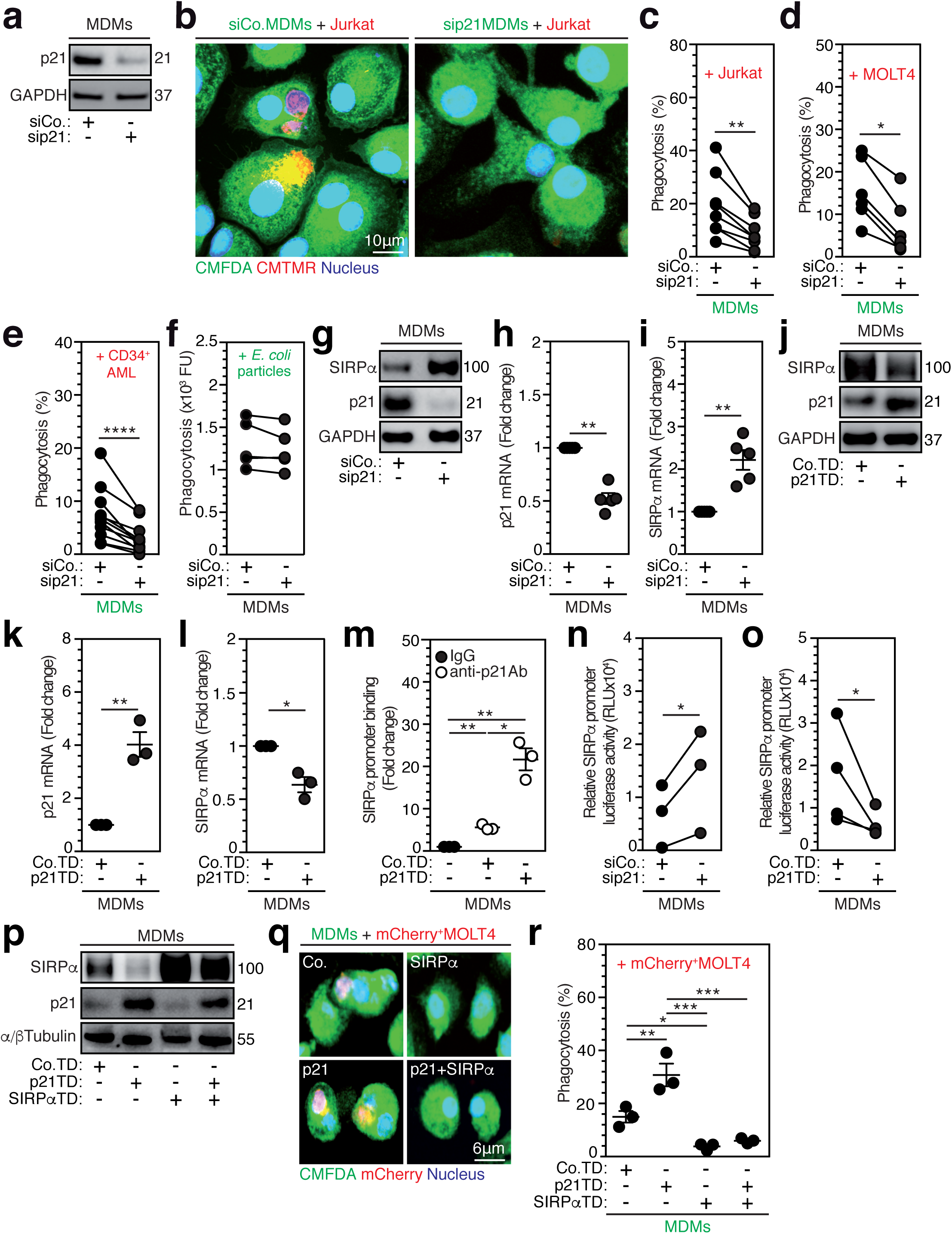
Macrophage-expressed p21 governs phagocytosis of leukemic cells through the repression of SIRPα transcription. **a,** p21 expression by WB in control (siCo.) or p21-silenced (sip21) MDMs after 24h siRNAs transfection. **b**, Confocal micrograph of siCo. or sip21 CMFDA^+^MDMs co-cultured with CMTMR^+^Jurkat cells for 8h. **c-e,** Phagocytosis percentages of Jurkat (**c**) MOLT4 (**d**) or patient AML CD34^+^ (**e**) cells by siCo. or sip21 MDMs after 8h co-culture (**p=0.0078, *p=0.0313, ****p<0.0001). **f**, Phagocytosis rate, determined by fluorescence units (FU), of pHrodo green *E. coli* bioparticles by siCo. or sip21 MDMs at 2h. **g-i,** SIRPα and p21 proteins by WB (**g**) or mRNAs by qPCR (**h,i**) in siCo. or sip21 MDMs after 24h silencing (**p=0.0024, **p=0.0020). **j-l,** SIRPα and p21 proteins by WB (**j**) or mRNAs by qPCR (**k,l**) in MDMs transduced with control (Co.TD) or p21 (p21TD) expressing lentiviral vectors after 72h transductions (**p=0.0063, *p=0.0291). **m**, ChIP-qPCR assays in control MDMs or in Co.TD (**p=0.0016) or p21TD (**p=0.0017, *p=0.0194) MDMs immunoprecipitated with control IgG or anti-p21 antibodies and qPCR detection of SIRPα promoter, after 72h transduction. **n**,**o**, Luciferase activities assessed at 24h after silencing in siCo or sip21 (**n**) or at 72h after transductions in Co.TD or p21TD (**o**) MDMs that were transduced with lentiviral vector expressing luciferase reporter gene under SIRPα promoter (*p=0.0431, *p=0.0458). **p-r,** MDMs derived from monocytes transduced with control (Co.TD) and/or p21 (p21TD) and/or SIRPα (SIRPαTD) expressing lentiviral vectors assessed for indicated proteins by WB (**p**) and for phagocytosis of mCherry^+^MOLT4 cells by confocal microscopy (**q**) and percentages (**r**) after 8h co-culture. In (**a**,**b**,**g**,**j**,**p**,**q)** data are representative of n=3 donors. In (**c**), (**d**), (**e)**, (**f**,**o**) and (**n**) data are donor-matched from n=8, n=6, n=9, n=4 and n=3 donors. In (**e**) CD34^+^ cells are from n=6 AML patients. In **(h**,**i)** and **(k**,**l**,**m,r)** data are means±SEM from n=5 and n=3 donors. *p<0.05, **p<0.01, ***p<0.001 and ****p<0.0001 are determined with two-tailed paired Wilcoxon (**c,d,e)**, two- (**h,i,k,m**) or one- (**l)** tailed ratio-paired t, one-tailed paired t (**n,o)** and ANOVA Tukey’s multiple comparison (**r**), tests.

To elucidate the molecular mechanisms promoting p21-dependent phagocytosis of leukemic cells, we evaluated simultaneously the expression of p21 and SIRPα, which is the main negative regulator of macrophage-mediated tumor cell phagocytosis^27^. Depletion of p21 in macrophages increased protein expression (Fig. 2g) and mRNAs of SIRPα (Fig. 2h,i), indicating that p21 down-regulates SIRPα protein through inhibition of its transcription. Reciprocally, up-regulation of p21 through transduction of MDMs with p21-expressing lentiviral vector (p21TD) (Fig. 2j) or with PMA treatment (Extended Data Fig. 3j) decreased the expression of SIRPα protein. Consistently, SIRPα mRNA was decreased in p21-transduced (p21TD) macrophages, compared with control-transduced (Co.TD) macrophages (Fig. 2k,l), further indicating that p21 represses the transcription of SIRPα.

Various mechanisms were shown to account for the ability of p21 to negatively regulate gene transcription^16,28,29^. Using chromatin immunoprecipitation (ChIP)-qPCR assays, p21 was detected at the level of SIRPα promoter in MDMs, and p21 overexpression was observed to significantly increase its binding to SIRPα promoter (Fig. 2m). When MDMs in which p21 expression had been decreased (Fig. 2n) or enforced (Fig. 2o) were transduced with a lentiviral vector expressing luciferase reporter gene under control of the endogenous macrophage *SIRPα* promoter, we observed, respectively, an increase or a decrease in *SIRPα* promoter-dependent luciferase activity, further demonstrating that p21 could repress *SIRPα* gene transcription. To evaluate the impact of the CDKN1A-SIRPα axis on leukemic cell phagocytosis, MOLT4 cells stably expressing fluorescent mCherry reporter gene (mCherry^+^MOLT4) were co-cultured with MDMs generated from genetically engineered human monocytes (Mo) transduced with an empty (Co.TD-Mo), a *CDKN1A*-expressing (p21TD-Mo), a *SIRPα*-expressing (SIRPαTD-Mo) lentiviral vector or combination of lentiviral vectors (p21+SIRPαTD-Mo). To overcome p21-mediated repression of *SIRPα*, we used a promoter, which was distinct from endogenous macrophage *SIRPα* promoter and did not respond to p21. As expected, enforced p21 expression decreased endogenous expression of SIRPα protein (Fig. 2p) and enhanced phagocytosis of mCherry^+^MOLT4 cells (Fig. 2q,r), while enforced expression of SIRPα (Fig. 2p) inhibited phagocytosis (Fig. 2q,r). Most importantly, transduction of a *SIRPα* cDNA that was insensitive to p21 repression abrogated phagocytosis of mCherry^+^MOLT4 cells by p21-overexpressing macrophages (Fig. 2p,q,r), demonstrating that p21-induced phagocytosis depends on SIRPα biological activity. p21 is unlikely to be involved in pro-inflammatory reprogramming of MDMs following leukemic cell engulfment, since p21 overexpression did not decrease cell surface expression of CD163 (Extended Data Fig. 3i), nor did it affect IRF5 expression (Extended Data Fig. 3j).

To explore how p21-mediated modulation of MDM phagocytic activity toward leukemic cells could be used therapeutically, we first showed that adoptive transfer of CFSE-labeled human monocytes in total body irradiated NOD.Cg-*Prkdc^scid^Il2rg^tm1Wjl^*/SzJ (NSG) mice could generate CFSE^+^ anti-inflammatory macrophages, detected in the bone marrow and the spleen within 7 days after monocyte adoptive transfer (Extended Data Fig. 4a-c,f). mCherry^+^MOLT4 cells were engrafted in NSG mice to develop a mouse model of human T ALL, as indicated by the presence of mCherry^+^ cells in the peripheral blood by day 21 after engraftment (Extended Data Fig. 4d,e and g), and body weight loss, bone marrow invasion and marked splenomegaly by day 35 after engraftment (Extended Data Fig. 4h,i). We then set up the adoptive transfer of CFSE^+^ p21TD-Mo or Co.TD-Mo, which did not induce any toxic effects (body and spleen weights; Extended Data Fig. 5a,b). p21TD-Mo or Co.TD-Mo-derived CFSE^+^ MDMs were equally distributed in the bone marrow and the spleen of NSG mice, without being detected in peripheral blood or liver (Extended Data Fig. 5c-e). We transferred genetically engineered CFSE^+^ p21TD-Mo or Co.TD-Mo into NSG mice that were simultaneously engrafted with MOLT4^+^mCherry cells (Fig. 3a). Interestingly, the adoptive transfer of p21TD-Mo prevented body weight loss (Fig. 3b,c) and splenomegaly development (Fig. 3d,e) in mCherry^+^MOLT4-engrafted NSG mice. These clinical parameters were associated with a significant reduction of the leukemic burden in the peripheral blood, the bone marrow, the spleen and the liver of p21TD-Mo-treated mice, compared with Co.TD-Mo mice (Fig. 3f-j and Extended Data Fig. 5f). Consistently, a significant increase in survival was also observed in p21TD-Mo-treated leukemic mice (Fig. 3k). Altogether, these results demonstrated that the adoptive transfer of p21TD-Mo strongly reduced progression of leukemia.

**Fig. 3.**
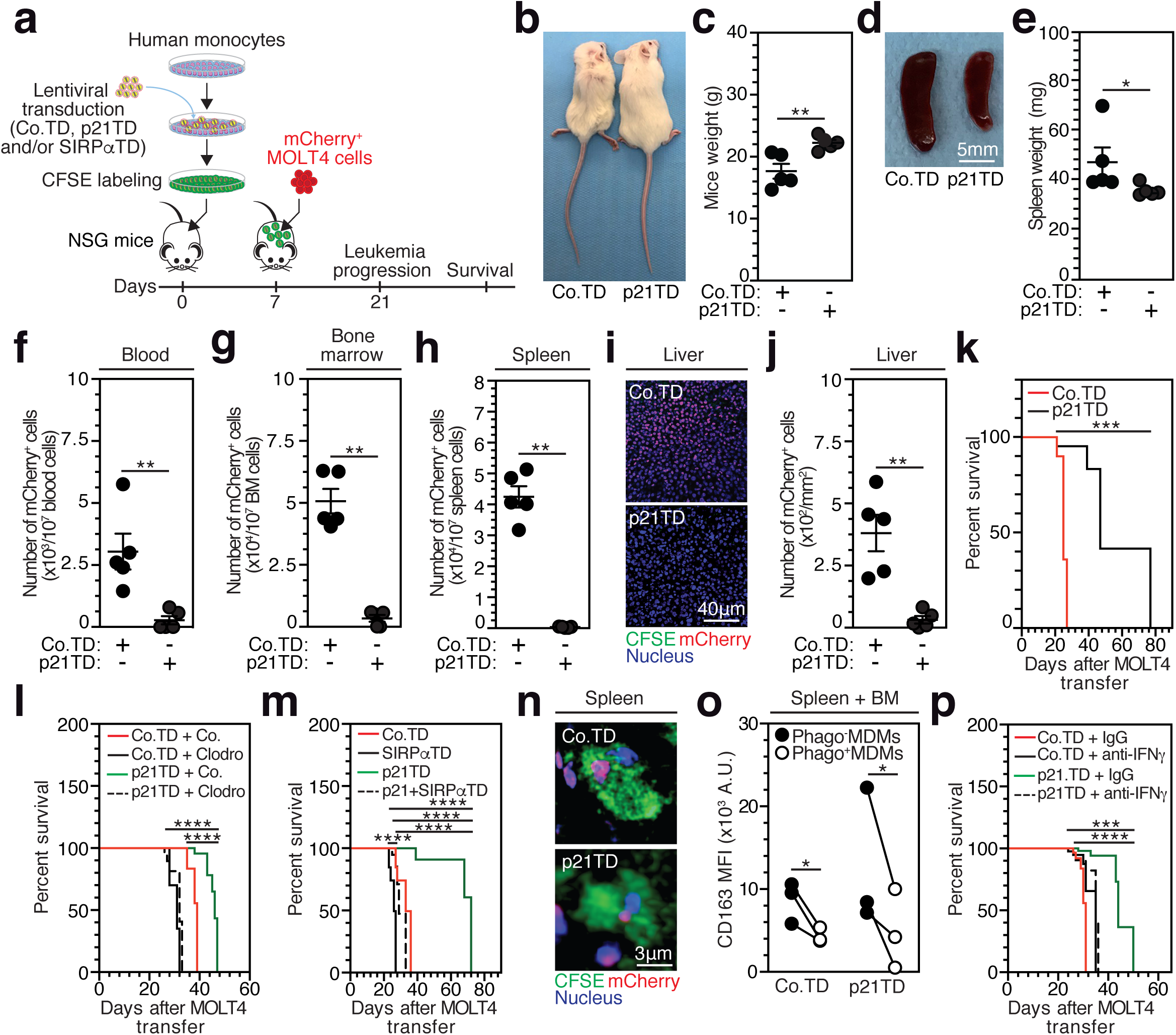
Prophylactic adoptive transfer of p21TD-Mo decreases leukemia burden and elongates mice survival in a model of human T-ALL. **a,** Schematic showing prophylactic adoptive transfer of CFSE-labeled, control (Co.TD), p21 (p21TD) and/or SIRPα (SIRPαTD) genetically engineered human monocytes (Mo) into total body irradiated (TBI) NSG mice which will be engrafted after 7d with mCherry^+^MOLT4 cells. **b-k,** Engrafted mice, which received indicated monocytes, were assessed at 21d for body (**p=0.0079) (**b**,**c**) and spleen, (*p=0.00317) weights (**d**,**e**), for leukemia burden in blood, bone marrow and spleen (**p=0.0079) by FACS (all **p=0.0079) (**f**,**g**,**h**) and in liver tissues by confocal microscopy (**p=0.0079) (**i**,**j**), and for survival (***p=0.0003) (**k**). **l,** Survival of engrafted mice, which received indicated monocytes, were treated 21d after monocyte transfers with control (Co.) or clodronate (Clodro) containing liposomes (****p<0.0001). **m,** Survival of engrafted mice that received indicated monocytes (****p<0.0001). **n,o,** Confocal micrographs of phagocytic (CFSE^+^mCherry^+^) macrophages in spleen (**n**) and CD163 membrane expression by FACS on sorted single CFSE^+^ (Phago^-^) MDMs or phagocytic CFSE^+^mCherry^+^ (Phago^+^) MDMs from spleen and bone marrow (*p=0.033)(**o**) from engrafted mice which received indicated monocytes. **p,** Survival of engrafted mice that received indicated monocytes and were treated 15d after monocyte transfers with isotype control (IgG) or with anti-IFNγ blocking antibodies (100 μg/mice) (***p=0.0002, ****p<0.0001). In (**b**,**d**,**i**,**n**) data are representative of n=5 mice/group. In (**c,e,f,g,h,j**) data are means±SEM from n=5 mice/group. In (**k**,**m**), (**l**) and (**p**) survival data are from n=5, n=6 and n=7 mice/group. In (**o**) data are mouse-matched from n=3 mice/group. *p<0.05, **p<0.01, ***p<0.001 and ****p<0.0001 are determined with two-tailed unpaired Mann-Whiteny (**c**,**e**,**f**,**g**,**h**,**j**), ANOVA Friedman (**o**) and Mantel-Cox (**k**,**l**,**m**,**p**), tests.

To decipher the biological mechanisms involved in the beneficial effect of p21TD-Mo, mCherry^+^MOLT4 engrafted NSG mice (obtained as shown in Fig. 3a) were treated 21 days after the adoptive transfer of p21TD-Mo or Co.TD-Mo with liposomes containing or not clodronate. After 24 hours of treatment, clodronate selectively depleted p21TD-Mo or Co.TD-Mo-derived CFSE^+^ macrophages (Extended Data Fig. 6a-c) and strongly reduced the survival of p21TD-Mo-treated NSG mice (Fig. 3l). These results further demonstrate the key role of p21TD-Mo-derived macrophages in the inhibition of leukemia progression. To address *in vivo* the impact of p21-mediated SIRPα repression on disease regression, p21TD-Mo, SIRPαTD-Mo, p21+SIRPαTD-Mo or Co.TD-Mo were engineered (as shown in Fig. 2p) and transferred into NSG mice that were engrafted with mCherry^+^ MOLT4 cells (Fig. 3a). As expected, while p21TD-Mo treated NSG mice lived longer, a significant decrease in survival of mice that received SIRPαTD-Mo was observed (Fig. 3m). In agreement with previous results (Fig. 2q,r), exogenous expression of a *SIRPα* cDNA that is resistant to p21 repression into p21TD-Mo abrogated the positive effect of engineered p21TD-Mo transfer on mouse survival (Fig. 3m), further demonstrating the role of p21-mediated SIRPα repression in regulation of MDM function. The decreased survival of mice after depletion of Co.TD-Mo-derived macrophages with clodronate-containing liposomes (Fig. 3l) and adoptive transfer of SIRPαTD-Mo (Fig. 3m) confirmed our *in vitro* results (Figs. 1 and 2), and demonstrated that lentiviral transduction of p21 potentiates the intrinsic capacity of anti-inflammatory macrophages to engulf leukemic cells *in vivo*.

Considering the pro-inflammatory fate of Phago^+^MDMs detected *in vitro* (Fig. 1), we sought to investigate its occurrence *in vivo* and its relevance for p21TD-Mo-based therapy. Twenty-one days after the adoptive transfer of engineered CFSE^+^ monocytes, CFSE^+^ Co.TD-Mo or p21TD-Mo-derived macrophages engulfing mCherry^+^MOLT4 cells were detected in the spleen of treated mice (Fig. 3n). More importantly, flow cytometry analysis of FACS-sorted non-phagocytic CFSE^+^ (Phago^-^) or phagocytic (Phago^+^) CFSE^+^mCherry^+^ MDMs from the spleen and the bone marrow of treated mice showed that Phago^+^MDMs decreased the membrane expression of CD163 scavenger receptor, compared with Phago^-^MDMs (Fig. 3o), thus confirming that *in vivo,* phagocytosis of leukemic cells promoted activation of MDMs toward a pro-inflammatory phenotype.

Given the key role of IFNγ secretion by Phago^+^MDMs in the *in vitro* pro-inflammatory reprogramming of bystander Phago^-^MDMs (Fig. 1), p21TD-Mo transferred NSG mice that were engrafted with mCherry^+^MOLT4 cells (Fig. 3a) were treated with anti-human IFNγ (hIFNγ) blocking antibodies 15 days after monocyte transfer and analyzed for survival. Treatment with anti-hIFNγ blocking antibodies strongly reversed the increased survival of mice that received p21TD-Mo (Fig. 3p), compared with controls. These results indicate that p21TD-Mo-based cellular therapy drives the engraftment of p21-transduced phagocytes, which besides directing the elimination of leukemic cells, triggers the secretion of the pro-inflammatory cytokine IFNγ and supports the pro-inflammatory reprogramming of macrophages, which in turn participates in regression of leukemia.

To further explore the therapeutic effect of p21TD-Mo based cellular therapy, we treated NSG mice engrafted with patient-derived T-ALL cells (Extended Data Table 3), including two patient-derived xenograft models derived from the same patient at diagnosis (PDX#1) and at relapse (PDX#2) (Fig. 4a and Extended Data Fig. 7a-f). The adoptive transfer of p21TD-Mo significantly increased the survival of T-ALL PDX engrafted mice, compared with control mice treated with Co.TD-Mo (Fig. 4b-d). Moreover, the adoptive transfer of p21TD-Mo-based cellular therapy remained effective in mice engrafted with T-ALL cells at relapse. Altogether, these results support the translational potential of the therapeutic induction of p21-mediated tumor phagocytosis for treatment of T ALL.

**Fig. 4.**
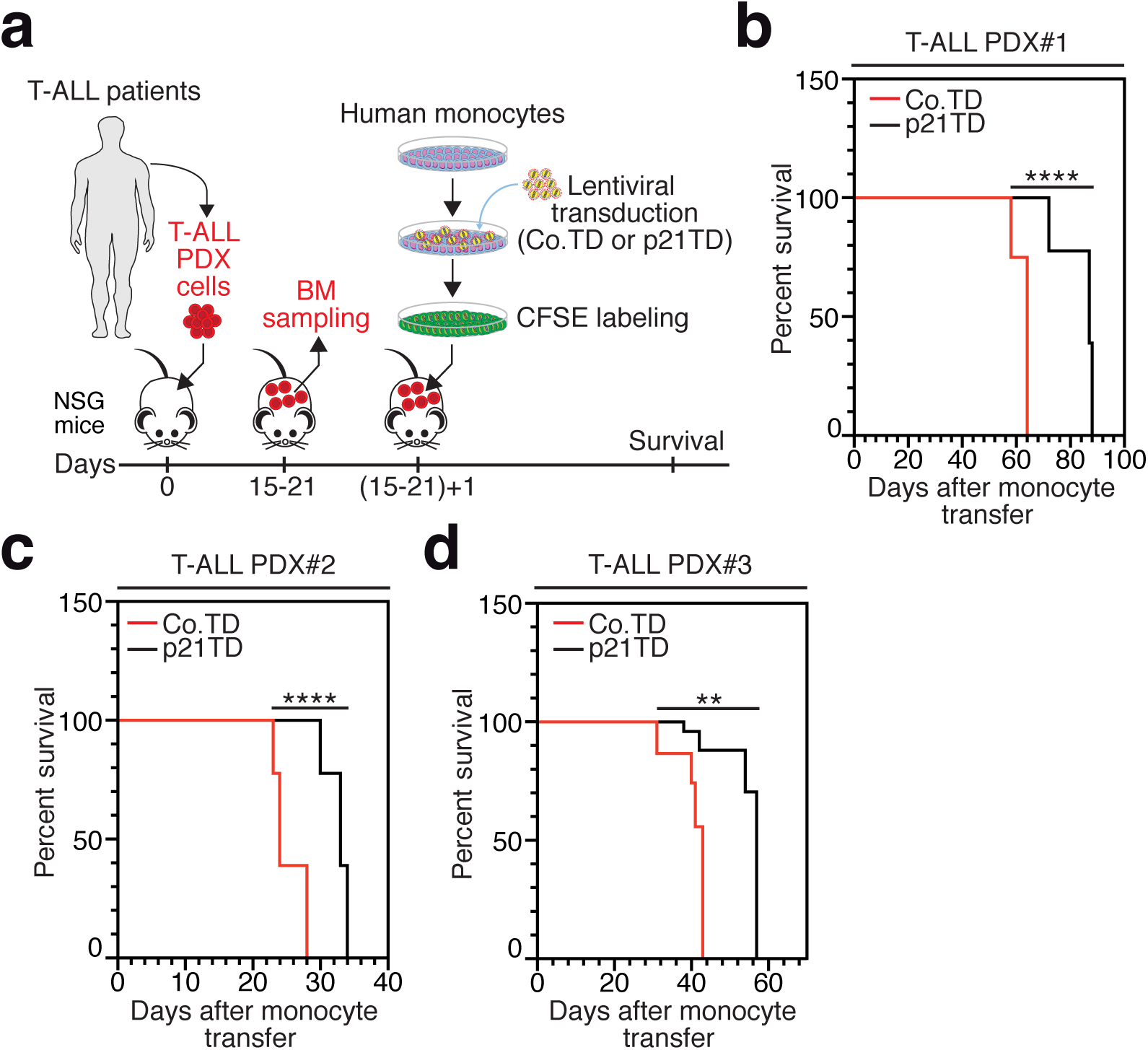
Curative adoptive transfer of p21TD-Mo elongates mice survival in diagnosed or relapsed T-ALL-derived PDX models. **a,** Schematic showing curative adoptive transfer of Co.TD-Mo or p21TD-Mo in NSG mice that were engrafted with T-ALL PDXs. **b-d,** Survivals of mice that were engrafted with T-ALL PDX#1 (****p<0.0001) (**b**), PDX#2 (****p<0.0001) (**c**) and PDX#3 (**p=0.003) (**d**) and received curatively, as shown in (**a**), Co.TD-Mo or p21TD-Mo. PDX#1 and PDX#2 are from the same patient isolated at diagnosis and relapsed stages, respectively. In (**b**-**d**) survival data are from n=5 mice/group. **p<0.01 and ****p<0.0001 are determined with Mantel-Cox (**b-d**) test.

## Discussion

Taken together, our findings demonstrated that macrophage-expressed p21 is responsible for the pro-inflammatory activation of macrophages through the induction of tumor phagocytosis. Our findings provide direct genetic and functional evidence that macrophage-expressed p21 is a key regulator of leukemic cell phagocytosis by repressing the transcription of the inhibitory receptor of phagocytosis, SIRPα. After degradation of engulfed cells, phagocytic macrophages undergo pro-inflammatory activation and, through the secretion of IFNγ, convert surrounding anti-inflammatory macrophages into pro-inflammatory cells. Supporting the translational potential of our findings, the adoptive transfer of p21TD-Mo into mouse models of human leukemia cells leads, after their differentiation into TAMs in the spleen and bone marrow of treated mice, to phagocytosis of leukemic cells and the pro-inflammatory reprogramming of engineered TAMs. The p21-initiated, SIRPα-repressed, phagocytosis-guided, IFNγ-dependent pro-inflammatory macrophage activation of TAMs strongly reduces leukemic burden and prolongs mouse survival. Our findings uncover a novel cellular mechanism to direct the killing of leukemic cells and harness anti-tumor innate immunity in a way that could be targeted for cellular therapy against leukemia. Our findings also identify SIRPα as a new target of transcriptional co-repressor p21^16,28,29^. Future studies will identify the upstream and downstream effectors of p21 that enable p21 binding to SIRPα promoter and contribute to the repression of SIRPα transcriptional activity. In addition to the well-demonstrated release of IFNγ by CD4^+^ T cells, NK cells and activated macrophages^30^, our results on the secretion of IFNγ by phagocytic macrophages after leukemic cell degradation supports the notion that the accumulation in the tumor microenvironment of p21TD-Mo-derived macrophages with enhanced abilities to engulf leukemic cells and to trigger *in situ* the pro-inflammatory reprogramming of TAMs, could be an attractive alternative to overcome tumor immune evasion strategies^3^. Our findings also support the hypothesis that, in syngeneic immunocompetent murine models of leukemia, the effectiveness of p21TD-Mo-based cellular therapy would be increased, since phagocytic macrophages in the tumor microenvironment would activate, besides bystander macrophages, other host immune cells to support anti-tumor T cell priming and favor an adaptive anti-tumor immune response.

The therapeutic feasibility and safety of autologous macrophage-based cell therapy was previously shown^31,32^. However, genetic engineering of primary myeloid cells with clinically approved vectors such as self-inactived human immunodeficiency virus 1 (HIV-1)-based lentivirus remains for a long time a major difficulty^32^. In this study, we circumvent the resistance of primary monocytes and macrophages to lentiviral transduction by co-transducing p21-expressing lentiviral vector with viral like particles containing Vpx protein, which was shown to degrade SAMHD1 viral restriction factor without affecting p21 expression^16^, and thus enables to evaluate the potential of p21TD-Mo-based cellular therapy. Adoptive transfer of p21TD-Mo would represent a broad-spectrum therapy for leukemia because it could enhance the macrophage capacity for phagocytosis without affecting antigen pressure selection, which could induce the emergence of clonal resistance^33^. Finally, considering the current difficulties in eradicating acute leukemia, the adoptive transfer of p21TD-Mo should be considered as a novel strategy that could complement commonly used chemotherapeutic approaches.

## Methods

### Primary cells and cell lines

Monocytes, MDMs and PBLs were obtained and differentiated as previously described^16,34^, from peripheral blood mononuclear cells (PBMCs) of buffy coats of healthy donors from French blood bank (Etablissement Français du Sang) in accordance with French law and with written informed consent of each donor. Monocytes were isolated from PBMCs by adherence to the plastic in macrophages medium (MM) (RPMI supplemented with 200 mM L-glutamine, 100 U/ml of penicillin, 100 μg/ml streptomycin, 10 mM HEPES, 10 mM sodium pyruvate, 50 μM β-mercaptoethanol, 1% minimum essential medium vitamins, 1% non-essential amino acids, all from Gibco) containing 2% (vol/vol) of heat inactivated (HI, for 1h at 56°C) human AB serum (hABS) (Sigma, #H3667). After extensive washings with DPBS (Gibco, #14190-094) to eliminate non-adherent cells, monocytes were incubated overnight in MM medium containing 10% HI hABS before in vitro differentiation or with 2% HI fetal bovine serum (FBS) before genetic engineering and/or adoptive transfer. Monocyte purity was analyzed by FACS and revealed that 90 to 96% cells expressed hCD11b and hCD14 and did not express hCD56 (NK cells), hCD3 (T cells) and hCD20 (B cells) markers. For differentiation into MDMs, monocytes were cultured for 6 to 7 days in hydrophobic Teflon dishes (Lumox, #94.6077.305) in MM containing 20% HI hABS yielding adherent non-proliferating cells expressing 90-96% macrophage (hCD11b, hCD14, hCD71) and M2-like (hCD163 and hCD206) markers. MDMs were then cultured in MM containing 10% of HI FBS at 1×10^6^/ml for experiments. Isolation, PHA-P/IL2 activation and culture of PBLs were performed as previously described^34^. Jurkat, MOLT4, CEM, HEL and K562 cells were all from American Type Culture Collection (ATCC) and cultured in RPMI with 10% HI FBS except for K562 (grown in IMDM (Gibco, #21980-032) with 10% HI FBS. For phagocytosis assays, all target cells were checked for freedom from mycoplasma contamination and were co-cultured with MDMs in MM with 10% HI FBS.

### Patient samples

Peripheral blood (PB) and bone marrow (BM) samples from AML patients (Extended Data Table 1) were prospectively collected after the obtention of informed consent according to the Declaration of Helsinki in the context of MYELOMONO2 study of Groupe Francophone des Myelodysplasies and approval by ethical committee authorizations (CCTIR N°14-266 and CNIL N°914283). Mononucleated cells from PB or BM were then enriched for CD34^+^ cells on the AutoMacs system using human CD34 MicroBead kit (Miltenyi Biotec, #130-046-702) according to manufacturer’s instructions. The CD34 cell purity was checked by FACS and cells were cultured in IMDM with 10% HI FBS supplemented with human cytokine and growth factor cocktail from Peprotech (hTPO (#300-18) (10 ng/ml), hSCF (#300-07) (25 ng/ml), hIL3 (#200-03) (10 ng/ml), hIL6 (#200-06) (10 ng/ml), hFlt-3 (#AF-300-19) (10 ng/ml)) for at least three days before phagocytosis assays. T-ALL PDXs samples were obtained as previously described^35^.

### Plasmids, lentiviral vectors and transductions

For genetic engineering of monocytes, MDMs and MOLT4 cells, self-inactivated (SIN) lentiviral vectors were used. p21 cDNA was cloned in pAIP lentiviral vector^36^ under SFFVp promoter and SIRPα cDNA was cloned in pRRL-EF1-PGK-GFP lentiviral vector under EF1 promoter. mCherry cDNA was cloned in pRRL lentiviral vector under EF1 promoter for the establishment of mCherry^+^MOLT4 stable cell line. SIRPA 2 promoter sequence (EP026655) was identified from Eukaryotic Promoter Database (EPD) to be expressed in human primary monocytes and macrophages, was fused to luciferase (Luc) reporter gene and cloned in pRRL-EF1-PGK-GFP lentiviral after the insertion of three stop codons at the 3’ end of E2F1 promoter. The initiator motif of SIRPA 2 promoter is at -530 bp before transcription starting site (TSS) of SIRPα gene. Controls (AIP or RRL-EF1-PGK-GFP) or gene-encoding (AIP-p21, RRL-EF1-SIRPα-PGK-GFP, RRL-EF1-3xstop codons-SIRPA 2-Luc-PGK-GFP or RRL-mCherry) lentiviral vectors were produced by transfecting 5×10^6^ HEK293T cells on T150 plate in 10 ml Opti-mem medium (Gibco, #31985-070) (containing 2% HI FBS and 100 U/ml penicillin, 100 μg/ml streptomycin) with 4 μg pDM2-VSV-G, 10 μg pΔ8.91 packaging and 8 μg lentiviral vector plasmid using Fugene (Promega, #E2312) according manufacturer’s instructions. Cell supernatants were removed after 24h transfection and replaced by fresh 10 ml Opti-mem medium (containing 2% HI FBS and 100 U/ml penicillin, 100 μg/ml streptomycin) for additional 48h. Then, cell supernatant was harvested and filtered through a 0.45 μm pore size filter. Viral stocks were quantified for viral CAp24 content by ELISA (Perkin Elmer, #NEK050A) and by qPCR quantification of lentiviral integrated copies in human genome of MDMs as previously described^34,37^. Viral like particles containing Vpx protein (VLPs-Vpx^+^) were produced, as described above for lentiviral vectors, by co-transfecting 5×10^6^ HEK293T cells with 4 μg pSIV3^+^(Vpx^+^) (kindly provided by A. Cimarelli, ENS Lyon) and 4 μg pDM2-VSV-G. Monocytes (10^7^) or MDM (10^6^) were treated for 1h at 37°C with 1 ml of VLPs-VPX^+^ containing 5 μg/ml polybrene (Sigma, #H9268) for SAMHD1 degradation^16,37^ before the addition of 20 µg CAp24 of each control or/and encoding-gene lentiviral vectors with 5 μg/ml polybrene for 2h transduction. After extensive washings with medium, monocytes were differentiated for 7d in MM with 20% HI hABS before WB analysis and phagocytosis assays. MDMs were cultured 72h before mRNA qPCR analysis, luciferase activity using luciferase assay system (Promega, #E1500) or chromatin immunoprecipitation (ChIP)-qPCR assays. While genetically engineered monocytes (Co.TD-Mo or p21TD-Mo) used for adoptive transfer in NSG mice were further incubated for 30 min with 1 μM CellTracer CFSE cell proliferation kit (Invitrogen, #C34554) before extensive washings with DPBS and intravenous (IV) injection of mice. MOLT4 cells (5×10^6^) were transduced with 100 μg CAp24 RRL-mCherry by 1h spinoculation at 1200g at 25 °C and 1h incubation at 37°C. After 72h, MOLT4 cells expressing mCherry gene were sorted by FACS and cultured in RPMI with 10% HI FBS.

### *In vitro* macrophage-mediated phagocytosis assays, sorting and functional assays with Phago^-^MDMs and Phago^+^MDMs

For phagocytosis assays, MDMs (0.125×10^6^ cells) were let to adhere for 2h in 125 μl MM with 10% HI FBS in one well of 8-chamber tissue culture treated glass slide (Falcon, #354118), labeled with 20 μM cell tracker green CMFDA (Invitrogen, #C2925) for 1h at 37°C and washed three times with 1 ml of medium before co-culture with target cells. Leukemic, PBL or CD34^+^AML cells were incubated at 1×10^6^/ml in RPMI with 10% HI FBS containing 100 μM ZVAD (Bachem, #4027403.0005) and 10 μM cell tracker orange CMTMR (Invitrogen, #C2927) for 1h at 37°C. After extensive washings, CMTMR^+^ target cells (0.125×10^6^ cells) were added to adherent CMFDA^+^MDMs (1:1 ratio) at 1×10^6^ cells/ml in MM with 10% HI FBS for 8h co-culture in the presence of ZVAD (100 μM), Y27632 (30 μM) (Tocris, #1254), human recombinant Annexin V (5 μg/ml) (Invitrogen, #BMS306) or with the same amounts of their respective solvent (DMSO or H2O) or IgG1 isotype controls (Abcam, #ab280974). MDMs were pre-treated with Y27632 (30 μM, 24h), PMA (30 ng/ml, 32h) (Sigma, #P1585), MS275 (1 μM, 32h) (Enzo, #ALX-270-378-M001) or with same volume amount of DMSO before co-cultures with target cells. MDMs were also let to adhere on wells precoated with 100 mg intravenous human immunoglobulin (IVIg) (CSL Behring) for 24h before co-culture. Silenced MDMs for p21 (sip21) and its control (siCo.) were co-cultured with target cells or with pHrodo green bacterial *E. coli* bioparticles (according manufacturer’s instructions) (Life technologies, #P35366) at 24h or 48h, respectively, after siRNA transfection. Jurkat or CD34^+^ AML cells were pretreated with 50 μM cisplatin (CDDP) (Mylan) for 24h to induce apoptosis before co-culture with MDMs.

After 8h of co-culture, the cell supernatant (which contains target cells that were not internalized by MDMs) was removed and MDMs were washed extensively and fixed 5 min with 2% PFA before confocal microscopy analysis. For the FACS sorting of Phago^-^MDMs and Phago^+^MDMs, MDMs (10×10^6^) resuspended at 1×10^6^/ml in MM with 10% HI FBS were labeled with 2.5 μM CMFDA for 1h at 37°C in 50 ml falcon tube. Target cells (MOLT4 or CD34^+^ AML) (10×10^6^) were resuspended at 1×10^6^/ml in RPMI with 10% HI FBS containing 100 μM ZVAD for staining with 1.25 μM CMTMR for 1h at 37°C in 50 ml falcon tube. After extensive washings with medium, MDMs and target cells (1:1 ratio) were co-cultured in 50 ml Falcon tube at 1×10^6^/ml in MM with 10% HI FBS for 2h at 37°C. Then, Phago^+^ (CMFDA^+^CMTMR^+^) MDMs and Phago^-^ (CMFDA^+^) MDMs were sorted by FACS. The purity of sorted populations was checked with FACS. Phago^+^MDMs and Phago^-^MDMs (at least 0.01×10^6^ each) were then let to adhere on wells of 8-chamber tissue culture treated glass slides for 2h and 96h before 2% PFA fixation and confocal microscopy analysis of target cell internalization by MDMs and their corresponding degradation, respectively. For gene expression microarray analysis, CD163 membrane expression by FACS, cell supernatant analysis by proteome profiler human cytokine panel A array (according manufacturer’s instructions) (RD, #ARY005), Phago^+^MDMs and Phago^-^MDMs (at least 0.1×10^6^ each) were cultured in MM with 10% HI FBS for 96h. For WB analysis, Phago^+^MDMs and Phago^-^ MDMs were cultured 7d. For study of bystander activation of macrophages, 2h after FACS sorting, Phago^-^MDMs (0.1×10^6^ in 1 ml of MM with 10% HI FBS) were adhered on 24 well plates in bottom chambers of Phago^-^MDMs or Phago^+^MDMs (0.1×10^6^ in 200 μl) adhered on 0.4 µm pore membranes of transwell cell culture inserts (Falcon, #353095) for 15d of co-culture in the presence of 1 μg/ml anti-hIFNγ antibody clone 25718 (RD, #MAB285) or mouse IgG2A control isotype (RD, #MAB003). Phago^-^MDMs at bottom chambers were then analysed by WB.

### Adoptive transfer of lentivirus-mediated genetically engineered human monocytes into human T-ALL mouse models

Mice studies were performed in accordance with protocols approved by the French Ethical Committee CEEA26 and following recommendations of proper use and care of animal experimentation. Mice experiments were performed using 6- to 8- weeks old female and male NOD.Cg-*Prkdc^scid^Il2rg^tm1Wjl^*/SzJ (NSG) immune-deficient mice purchased from Charles River Laboratories, maintained in specific pathogen-free conditions and randomized for homogenous mice body weight groups (20g to 23g) before experiments. Intravenous (IV) injections were performed with 200 μl cell suspension in DPBS/mouse via retro-orbital sinus under isoflurane gas anesthesia. For the assessment of the *in vivo* differentiation of human monocytes into macrophages, monocytes (Mo) (5×10^6^/mouse) were labeled with 1 μM CFSE for 30 min at 37°C, extensively washed and IV injected into NSG mice 24h after their 1 Gray (1 Gy) total body irradiation (TBI) with X-RAD-320 irradiator (Precision X Ray). Seven days after monocyte transfer, NSG mice were sacrificed and sorted CFSE^+^ cells from dissociated bone marrow (BM) and spleen cells were analyzed for human M2 like macrophage markers (hCD11b, hCD14 and hCD163). For the establishment of the mouse model of human T-ALL, FACS-sorted mCherry^+^MOLT4 cells (10^6^/mouse), obtained by RRL-mCherry lentiviral vector transduction as described above, were IV injected in NSG mice and then after 30d, mCherry^+^ leukemic cells which engrafted BM were sorted by FACS and cultured in RPMI with 10% HI FBS for *in vitro* macrophage-mediated phagocytosis assays and mice injections. To characterize leukemia progression, mCherry^+^MOLT4 cells (10^6^/mouse) were IV injected in 1 Gy TBI NSG mice 7d after irradiation. The percentage of mCD45^-^hCD45^+^mCherry^+^ cells detected in the peripheral blood (PB) was determined every week. After 35d, mice were sacrificed and the presence of leukemic cells in BM and spleen were analyzed by FACS. To characterize the engraftment of genetically engineered monocytes into mice, CFSE-labeled Co.TD-Mo or p21TD-Mo (5×10^6^/mouse) were IV injected in 1 Gy TBI NSG mice and sacrificed after 21d. The presence of CFSE^+^ derived macrophages in PB, BM, spleen and liver were determined by FACS and confocal microscopy. For the prophylactic adoptive transfer of monocytes, 1 Gy TBI NSG mice were IV injected with CFSE-labeled Co.TD-Mo or p21TD-Mo (5×10^6^/mouse) and after 7d were IV injected with mCherry^+^MOLT4 cells (10^6^/mouse). Mice were then analyzed for overall survival or sacrificed at 21d to determine leukemia burden (in PB, BM, spleen and liver by FACS and confocal microscopy). *In vivo* pro-inflammatory activation of engineered macrophages was also determined by sorting phagocytic (CFSE^+^mCherry^+^cells) and single (CFSE^+^) Co.TD or p21TD-derived macrophages from BM and spleen cells and analyzing hCD163 membrane expression by FACS. To determine impacts of Co.TD-Mo or p21TD-Mo-derived macrophages and their related hIFNγ secretions on the overall survival of mice, treated NSG mice were also IV injected with of 200 μl/mouse of clodronate- or control PBS-containing liposomes (Liposoma, #CP-005-005) or intraperitoneally injected with 100 μg/mouse IgG control isotype (RD, #MAB003) or anti-hIFNγ antibody (clone 25718; RD, #MAB285), 21d or 15d after monocyte transfer, respectively. Clodronate-mediated depletion of CFSE-labeled Co.TD-Mo or p21TD-Mo-derived macrophages was assessed by FACS 24h after treatments. To study effects of p21-mediated SIRPα repression on the overall survival of engrafted mice, prophylactic monocyte adoptive transfer (5×10^6^/mouse) was performed with genetically engineered Co.TD-Mo, p21TD-Mo, SIRPαTD-Mo or p21+SIRPαTD-Mo, which were transduced as described above with respectively equal amounts of AIP+RRL-PGk-GFP, AIP-p21+RRL-PGK-GFP, AIP+RRL-SIRPα-PGK-GFP or AIP-p21+RRL-SIRPα-PGK-GFP lentiviral vectors. For curative approach, T-ALL PDX cells (10^5^/mouse for PDX#1 and PDX#2 and 2×10^5^/mouse for PDX#3) were IV injected in NSG mice. PDX cell engraftments were assessed at 21d (for PDX#1 and PDX#2) and 15d (for PDX#3) by femoral BM sampling as previously described^35^ and FACS detection of hCD45^+^hCD7^+^ PDX cells. Then 1d after BM sampling showing 40 to 70% PDX cell engraftment in BM, NSG mice were randomized to have homogenous groups for PDX engraftments, IV injected with Co.TD-Mo or p21TD-Mo (5×10^6^/mouse) and monitored for overall survival.

### Human macrophage knockdowns

MDM knockdowns were performed as we previously described^16,34^ using 50 nM of non-targeting pool control siRNAs (UGGUUUACAUGUCGA CUAA, UGGUUUACAUGUUGUGUGA, UGGUUUACAUGUUUUCUGA, UGGUUU ACAUGUUUUCCUA) or p21 smart pool selected siRNA (AGACCAGCAUGACAGAUUU), all purchased from Dharmacon. MDMs were silenced for p21 for 24h before WB or co-cultures with target cells.

### Gene expression

Determination of p21 and SIRPα mRNAs by RT-qPCR were performed as we previously described^16,38^ using TaqMan gene expression predesigned probes for SIRPα (Hs00757426 S1), p21 (Hs00355782 m1) and GAPDH (Hs02758991 g1). Gene expression analysis of Phago^-^MDMs and Phago^+^MDMs were performed with Agilent® SurePrint G3 Human GE 8×60K Microarray (Agilent Technologies, AMADID 28004) with two technical replicates for each sample. Data are determined for each analyzed donor by the mean of two technical replicates of Log2 fold change (FC) of gene intensity ratio of Phago^+^MDMs/Phago^-^ MDMs. To produce a comprehensive analysis of modulated genes for their ability to regulate anti-inflammatory or pro-inflammatory processes or to be modulated by IFNγ-dependent signaling pathways or treatments, a systematic literature search with Pubmed science web was performed using key words “the gene name AND macrophages” and “ the gene name AND interferon gamma” (Extended Data Table 2).

### Chromatin immunoprecipitation (ChIP)-qPCR assays

Chromatin immunoprecipitation assays were performed with MDMs using Chip assay kit (Millipore, #17-295) and according to manufacturer’s instructions. Briefly, cross-linked cell chromatins were sheared by sonication during 10 min at 40 W (duty factor: 20%; peak incident power: 200; cycles per burst: 200) by Covaris S220 (Woodingdean, UK). Chromatins were then immunoprecipitated with 20 μg of anti-p21 Waf1/Cip1 (12D1) antibodies (Cell signaling, #2947) or with equal amount of rabbit IgG control isotype. DNA-bound to chromatin immunoprecipitates were finally eluted and analyzed by qPCR using Power SYBER Green PCR Master Mix (Applied Biosystems#4367659) for the detection of SIRPα promoter sequence (SIRPA 2) with specific primers SIRPA 2 forward (5’CCACCGAGACACCTGGCCAG3’) and SIRPA 2 reverse (5’AAGTGAACGCAGGGGGAAGG3’). Specificity of promoter sequence detection was checked by negative qPCR controls which target unrelated promoter sequence at -2000 bp upstream of 5’ end of SIRPA 2 promoter using Untarget-SIRPA 2 forward (5’CCGTGGGTCTCAATGGCTTC3’) and Untarget-SIRPA 2 reverse (5’GGGGGATTAGGAAACTGGAG3’) primers. The normalization of DNA amount was performed by quantifying albumin gene copies with qPCR using human genomic DNA (20 ng/µl) standard (Roche, #11691112001) as we previously described^16^.

### Western blot

WBs were performed as we previously described^16,38^ using anti-p21 Waf1/Cip1 (12D1) (#2947), anti-Myosin Light Chain 2 (MLC2) (#3672), anti-Phospho (Ser19) Myosin light chain 2 (#3671), anti-αTubulin (#3873) and anti-α/β-Tubulin (#2148) antibodies from Cell Signaling, anti-IL8 (#ab106350), anti-IRF5 (#ab21689), anti-ILb (#ab2105) and anti-iNOS (#ab3523) antibodies from Abcam, anti-IL6 antibody (RD, #AB-206-NA), anti-SIRPα antibody (Invitrogen, #PA1-30537) and anti-GAPDH antibody (Millipore, #MAB374).

### Flow cytometry

Cell samples were analyzed by FACS using BD LSR2 or BD LSRFortessa (BD Biosciences) and sorted using BD FACSAria III (BD Biosciences). Cell stainings were performed in DPBS containing 2% (for cultured cells) or 5% (for mouse cells) HI FBS at 4°C for 2h incubation with 1:100 antibody dilution. For blood samples, red cells were lysed with ACK buffer before incubations with antibodies and/or FACS analysis for mCherry^+^ or CFSE^+^ cells. Mouse cell samples were filtered through 100 μm pore size membrane filters before FACS staining and analyses. Antibodies used were anti-hCD14 clone 61D3 (eBioscience, #12-0149-109 42), anti-hCD11b clone ICRF44 (#301342), anti-hCD34 Clone 561 (#343606), anti-hCD7 Clone CD7-6B7 (#343120) from Biolegend and anti-hCD163 clone GHI/61 (#562669), anti-hCD71 (#555537), anti-hCD206 (#551135), anti-hCD56 clone B159 (#560916), anti-hCD3 (#555339), anti-hCD20 (#559776), anti-mCD45 clone 30-F11 (#560501), anti-hCD45 clone HI30 (#555485) from BD Pharmingen. Membrane expression of CD163 was determined by mean fluorescence intensities (MFI).

### Confocal microscopy

After fixation, co-cultures containing single CMFDA^+^MDMs and phagocytic CMFDA^+^CMTMR^+^MDMs were permeabilized with 0.1% triton in DPBS for 5 min at room temperature, extensively washed, incubated with DPBS containing 10% HI FBS and 1:1000 Hoechst 33342 (Invitrogen, #H3570) for nuclear staining and mounted onto microscope cover slips with Fluoromount G (Southern Biotech, #0100-01). Immunofluorescence staining with anti-LAMP2 antibody (Santa Cruz, #sc-18822) and acquisitions by confocal microscopy (SP8, Leica) of fixed stained cells were performed as we previously described^34^. Percentages of phagocytosis were determined by dividing number of CMFDA^+^CMTMR^+^MDMs on total number of MDMs quantified on at least 5 fields containing at least 100 MDMs of each glass well. For determination of cell surfaces and volumes, target cells were labeled with CMTMR at 10 µM and 1:1000 Hoechst 33342 for 30 min at 37°C, and live imaged in ibidi 8 well chamber slides (ibiTreat, #80826) by confocal microscopy (SP8, Leica) with z-stacks (0.22 µm step) acquisitions covering from the top to the bottom of cells. After 3D constructions, cell volumes and surfaces were measured by Volocity software (Quorum Technologies). For detection of CFSE^+^ and mCherry^+^ cells engrafted in mice tissues, fresh intact pieces of spleen, liver or longitudinal cut femur BM were incubated in 500 µl HI FBS containing 0.3% triton and 1:250 Hoechst 33342 for 12h at 4°C. Medium was then replaced with cold HI FBS with 1:250 Hoechst 33342 and conserved on ice up to confocal microscopy acquisition of each organ tissue in ibidi 8 well chamber slides. Mouse tissues (at least 2 mm^2^/organ) were imaged by confocal microscopy (SP8, Leica) using 40x oil objective, hybrid detectors (pinhole airy: 0.6; pixel size: 284 nm, surface of each image field surface is 290.62×290.62 µm^2^) at optimal optical sectioning (OOS) of 1 µm. Organ cell layers were then 3D constructed by Imaris 5.7 software (Bitplane AG), and CFSE^+^ and mCherry^+^ cells were quantified and normalized to 10 mm^2^ of tissue surface.

### Statistics

Statistical analysis was performed in GraphPad Prism 6.0 (GraphPad). Statistical tests, adjustments used for multiple comparisons, exact calculated p-values and n size of each figure are indicated in figure legends. For all figures, statistical significances were given as *p <0.05 and **p <0.01, ***p<0.001, ****p<0.0001.

## Acknowledgements

We gratefully acknowledge Prof. D. M. Ojcius for helpful scientific discussions, NH Theraguix for supporting our projects against cancer, and M. Morabito and L. Bencheikh for their technical support. This work has benefited from the facilities and expertise of the Imaging and Cytometry Platform (S. Salome-Desnoulez and T. Manoliu), Genomic and Bioinformatics Platforms (N. Droin and G. Meurice), UMS 3655 CNRS / US 23 INSERM. Gustave Roussy Cancer Campus Villejuif, France. This work was supported by funds from Agence Nationale de la Recherche (ANR-10-IBHU-0001, ANR-10-LABX33, ANR-11-IDEX-003-01 and ANR Flash COVID-19 “MacCOV”), Fondation Gustave Roussy, Institut National du Cancer (INCA 9414), The SIRIC Stratified Oncology Cell DNA Repair and Tumor Immune Elimination (SOCRATE), Care network (directed by X. Mariette, Kremlin Bicêtre AP-HP) and Université Paris-Saclay (to J.-L. Perfettini) and The SIRIC Stratified Oncology Cell DNA Repair and Tumor Immune Elimination 2.0 (SOCRATE 2.0, INCA-DGOS-INSERM 12551) (to A. Allouch).

## Author contributions

J.-L.P. conducted the study. A.A. and J.-L.P. designed the study. A.A., L.V., Y.Z., S.Q.R. and Y.L. performed experiments. J.C., D.S.-B., S.DB., E. S., F.L. and F.P. provided PDXs and AML samples. A.A., Y.L., F.P. and J.-L.P. analyzed the results. A.A. assembled the figures. A.A. and J.-L.P. wrote the initial draft. A.A., L.V., Y.Z., S.Q.R., Y.L. and J.C., D.S.-B., S.DB., E. S., F.L., F.P. and J.-L. P. provided advice and edited the initial draft.

## Materials & Correspondence

Correspondence and requests for materials should be addressed to J.-L.P.

