## Extended Data Figures and Tables for "CDKN1A is a target for phagocytosis-mediated cellular immunotherapy in acute leukemia"

**Extended Data Figure legends**

**Extended Data Fig. 1 Characterization of macrophage phagocytosis of leukemic cells**. **a,** Confocal micrographs of CMFDA-labeled MDMs and CMTMR-labeled Jurkat cells that were co-cultured for 8h and stained for LAMP2 lysosome marker. **b,** Expressions of Myosin Light Chain 2 (MLC2) and its phosphorylated/activated form (MLC2S19*) by WB in MDMs treated for 24h in absence or presence of Y27632 (30 μM) ROCK pathway inhibitor. **c,** Percentages of phagocytosis of Jurkat cells by control or Y27632-pretreated MDMs, as in (**b**), at 8h of co-culture in presence or absence of Y27632 (30 μM). **d,e,** Percentages of phagocytosis of viable or apoptotic (pretreated for 24h with CDDP (50 μM)) Jurkat (*p=0.05) (**d**) or patient AML $CD34^+$ (*p=0.0331) (**e**) cells by MDMs in presence or absence of human recombinant AnnexinV (5 μg/ml) for 8h of co-culture. **f,g,** Percentages of phagocytosis of indicated leukemic cells or non-transformed PBLs by MDMs in function of target cell volumes (**f**) and surfaces (**g**), both determined after 3D reconstruction by Volocity software from confocal microscopy 0.22 μm acquired z stacks. In (**a,d**) data are representative n=3 donors. In (**b,e**) and (**c**) data are donor-matched from n=3 and n=2 donors. In (**f,g**) data are means±SEM from n=3 donors and for target cells n=10 independent experiments. *p<0.05 was determined with ordinary two-way ANOVA (**b,c**) and one-tailed Mann-Whitney (**e**), tests.

**Extended Data Fig. 2 Pro-inflammatory activation of phagocytic macrophages**. **a,b,** Secretions of indicated pro-inflammatory cytokines determined by cytokine array in the cell supernatants of FACS sorted $Phago^+$MDMs or $Phago^-$MDMs, as in (**Fig. 1e**) that were obtained after 2h of co-culture of MDMs with MOLT4 (*p=0.0143, *p=0.0143, *p=0.05, *p=0.0143, *p=0.0143, *p=0.03) (**a**) or with $CD34^+$ blasts (**b**) and 96h after FACS sorting. In (**a**) and (**b**) data are donor-matched from (n=3 for Gro-α and IL27 n=4 for IL8, IL23, IL1-ra, MIF) and (n=2) donors. *p<0.05 is determined with one-tailed unpaired Mann-Whitney (**a**) test.

**Extended Data Fig. 3 p21 is a key regulator of the phagocytosis of leukemic cells without affecting macrophage activation**.

**a,b,** Expression of p21 by WB in controls or PMA (30 ng/ml) (**a**) or MS275 (1 μM) (**b**)-treated MDMs for 32h. **c,d,e,** Percentage of phagocytosis of Jurkat (**c**) or MOLT4 (**d,e**) cells by controls or PMA (*p=0.0342,**p=0.0079) (**c,d**) or MS275 (***p=0.0007) (**e**)-pretreated MDMs detected after 8h of co-culture. **f,** Expression of p21 by WB in control or intravenous immunoglobulin (IVIg)-stimulated MDMs for 24h. **g,** Percentage of phagocytosis of Jurkat cells by control or IVIg prestimulated MDMs detected after 8h of co-culture (*p=0.0439). **h,** Expressions of SIRPα and p21 by WB in control or PMA-treated MDMs for 32h. **i,** CD163 membrane expression by FACS of MDMs derived from Co.TD or p21TD genetically engineered monocytes and differentiated into MDMs for 7d, as in (**Fig. 2p**), and then assessed by FACS 4d later. **j,** Expressions of IRF5 and p21 by WB in control (siCo.) or p21-silenced (sip21) MDMs 24h after siRNAs transfection. In (**a**,**b**,**f**,**h**,**j**) data are representative of n=3 donors. In (**c,d,e,g,i,j**) data are donor-matched from n=3 donors. *p<0.05, **p<0.01, ***p<0.001 are determined with one-tailed ratio-paired t (**c,d,e**) and ANOVA Tukey's multiple comparison (**g**), tests.

**Extended Data Fig. 4 Establishment of the adoptive transfer of human monocytes and a mouse model for human T-ALL**. **a,** Schematic of the adoptive transfer of CFSE-labeled human monocytes, through IV injections, into TBI NSG mice, sorting and immune-phenotyping 7d after monocyte transfer of $CFSE^+$ cells obtained from bone marrow (BM) and spleen for human macrophage markers. **b,c,** FACS dot plots of sorted $CFSE^+$ cells expressing human leukocyte marker CD45 (hCD45) (**b**) from bone marrow and spleen of engrafted mice and expression of indicated human macrophage markers (**c**). **d,** Schematic of the engraftment of $mCherry^+$MOLT4 cells into TBI NSG mice of $mCherry^+$MOLT4 cells, weekly peripheral blood sampling and FACS detection of leukemic cells. **e,** FACS dot plots showing the detection of $mCherry^+$MOLT4 cells in peripheral blood of engrafted mice expressing hCD45, but not murine CD45 (mCD45) cells, 35d after leukemic cell injection. **f,** Percentages of sorted $hCD45^+CFSE^+$ cells expressing indicated macrophage markers obtained in (**b,c**). **g,** Percentages of $mCherry^+$MOLT4 cells detected at indicated time points in the peripheral blood of engrafted mice as in (**d**). **h,** Photographs of NSG mice engrafted (+) or not (-) with $mCherry^+$MOLT4 cells, as in (**d**), and their corresponding BM and spleen 35d after leukemic cell injection. **i,** Percentages of $mCherry^+$MOLT4 cells in BM and spleen of engrafted NSG mice are indicated. In (**b,c,e,h**) data are representative of n=3 mice. In (**f,j**) data are means±SEM from n=3 mice. In **(g)** data are mouse-matched from n=3 mice.

**Extended Data Fig. 5 Biological parameters associated with the p21TD-Mo-based therapy for human T-ALL. a,b,** Body (**a**) and spleen (**b**) weights of NSG mice that were infused with CFSE-labeled engineered monocytes (Mo) (Co.TD-Mo or p21TD-Mo) or controls (Co.). Weights were determined 21d after monocyte IV injections. **c,** Numbers of $CFSE^+$ cells detected for $10^7$ cells of peripheral blood, bone marrow (BM) or spleen obtained from mice (that were infused with CFSE-labeled Co.TD-Mo or p21TD-Mo) were determined by FACS at 21d after monocytes injections. **d,** Confocal micrographs showing 21d after monocyte transfer, $CFSE^+$ cells in the spleen of mice that were infused with CFSE-labeled Co.TD-Mo or p21TD-Mo. **e,** Numbers of $CFSE^+$ cells in 1 $mm^2$ of bone marrow, spleen or liver tissues obtained from mice that were infused with CFSE-labeled Co.TD-Mo or p21TD-

Mo, and imaged by confocal microscopy 21d after monocyte injection. **f,** Confocal micrographs showing mCherry$^+$MOLT4 cells and CFSE$^+$cells in the spleen of engrafted NSG mice 21d after monocyte transfer, as in (**Fig. 3a**). In (**d**,**f**) data are representative of n=5 mice/group. In (**a,b**) data are means±SEM from n=3 mice for Co. group and n=5 mice for Co.TD or p21TD group. In (**c,e**) data are means±SEM from n=5 mice mice/group.

**Extended Data Fig. 6 Clodronate containing liposomes-mediated depletion of CFSE$^+$ Co.TD-Mo and p21TD-Mo-derived macrophages. a**, FACS dot plots of spleen CFSE$^+$ cells obtained from spleen of p21TD-Mo transferred NSG mice, as in (**Fig. 3a**)**,** which were treated with control (Co.) or clodronate (Clodro) containing liposomes 21d after monocyte transfer and assessed for specific depletion by FACS 24h after treatments. **b,c,** Numbers of CFSE$^+$ cells detected for $10^7$ cells of spleen (**b**) or bone marrow (BM) (**c**) obtained from mice which received Co.TD-Mo or p21TD-Mo, as shown in (**Fig. 3a**)**,** and were treated with Co. or Clodro containing liposomes 21d after monocyte transfer. The depletion of CFSE$^+$ cells was determined by FACS 24h after liposomal treatments. In (**a**) data are representative of n=3 mice/group. In **(b,c)** data are means±SEM from n=3 mice/group. ***p<0.001 and ****p<0.0001 are determined with ANOVA Tukey's multiple comparison (**b,c**) test.

**Extended Data Fig. 7 Characterization of the bone marrow invasion of mice which were engrafted with diagnosed or relapsed T-ALL-derived PDXs. a,bc,** FACS dot plots showing the bone marrow invasion of mice which were engrafted with T-ALL PDX#1 (**a**), PDX#2 (**b**) and PDX#3 (**c**), as shown in (**Fig. 4a**). PDX#1 and PDX#2 are from the same patient isolated at diagnosis and relapsed stages, respectively. The bone marrow invasion was assessed by FACS by detecting human leukocyte (hCD45) and T-ALL (hCD7) markers 24h before the adoptive transfer of Co.TD-Mo or p21TD-Mo. **d,e,f,** Percentages of hCD45$^+$hCD7$^+$

cells detected in the bone marrow of mice which were engrafted with T-ALL PDX#1 (**d**), PDX#2 (**e**) or PDX#3 (**f**), as shown in (**Fig. 4a**), were determined by FACS 24h before the adoptive transfer of Co.TD-Mo or p21TD-Mo. In (**a,b,c**) data are representative of n=10 mice. In (**d,e,f)** data are means±SEM from n=5 mice/group.

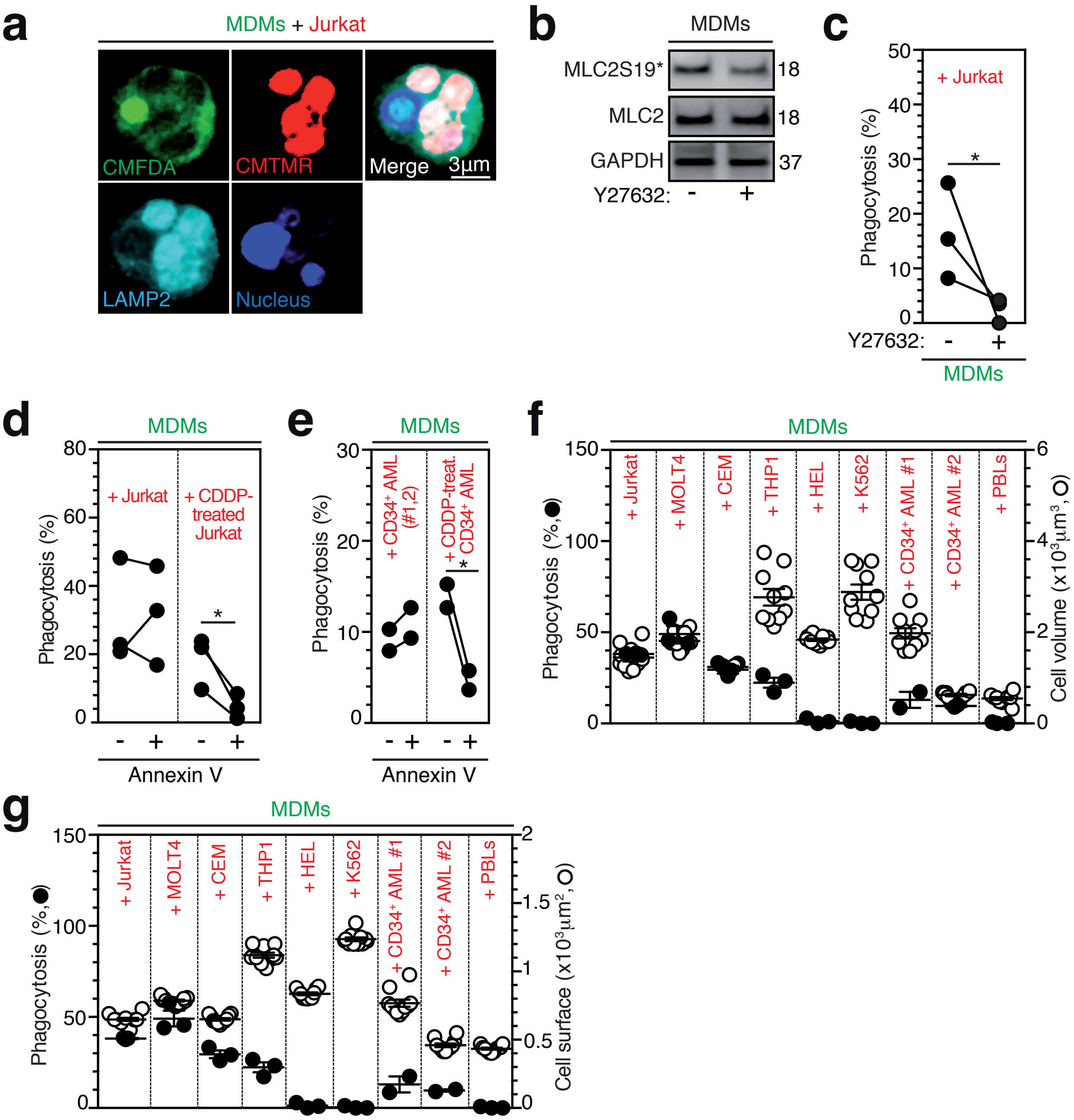


### ALLOUCH#EXTENDED1

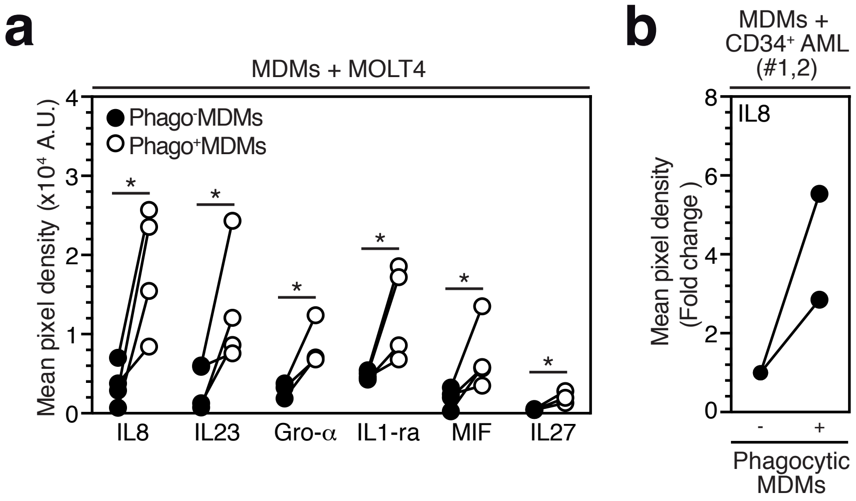


### ALLOUCH#EXTENDED2

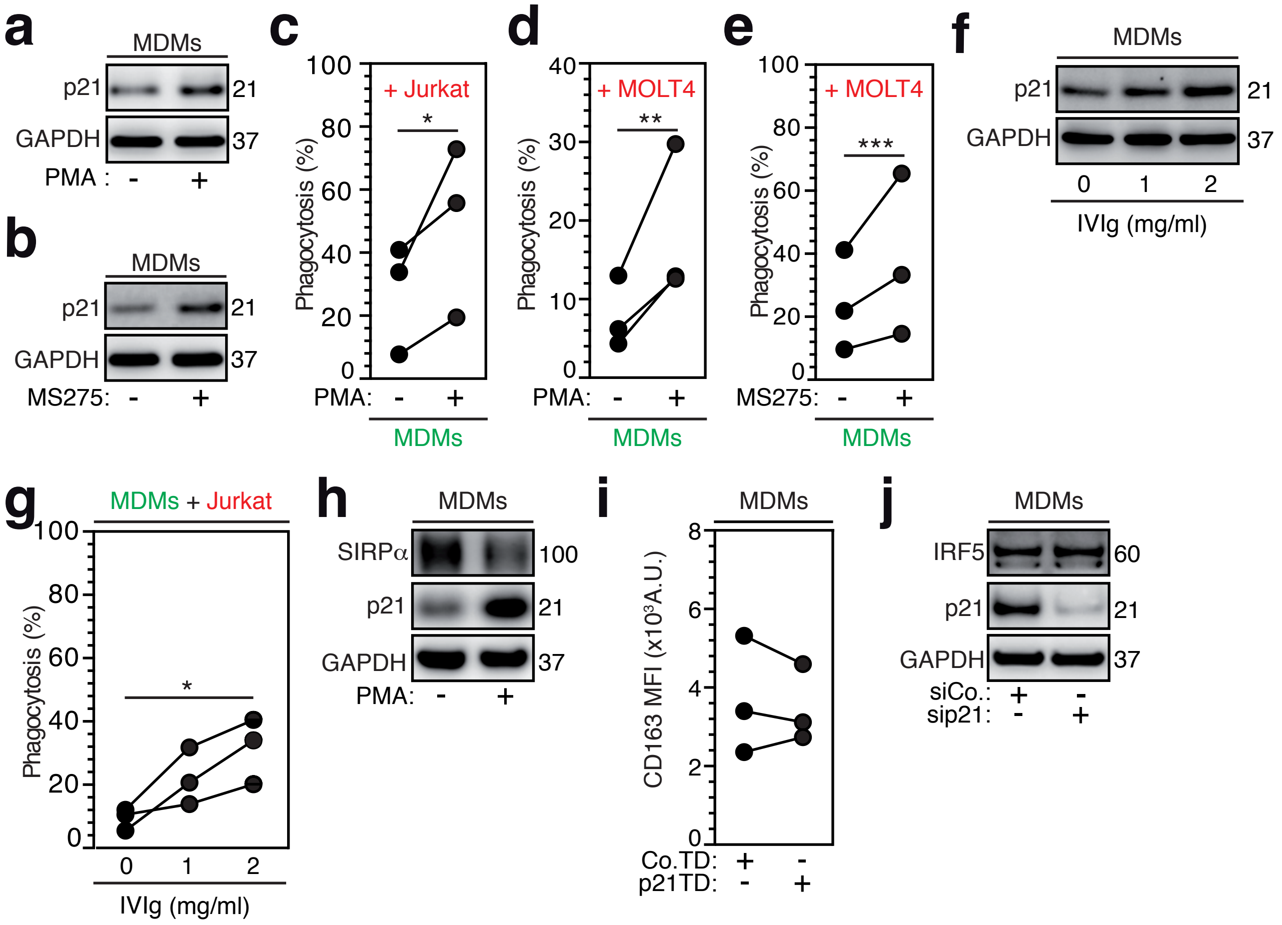


### ALLOUCH#EXTENDED3

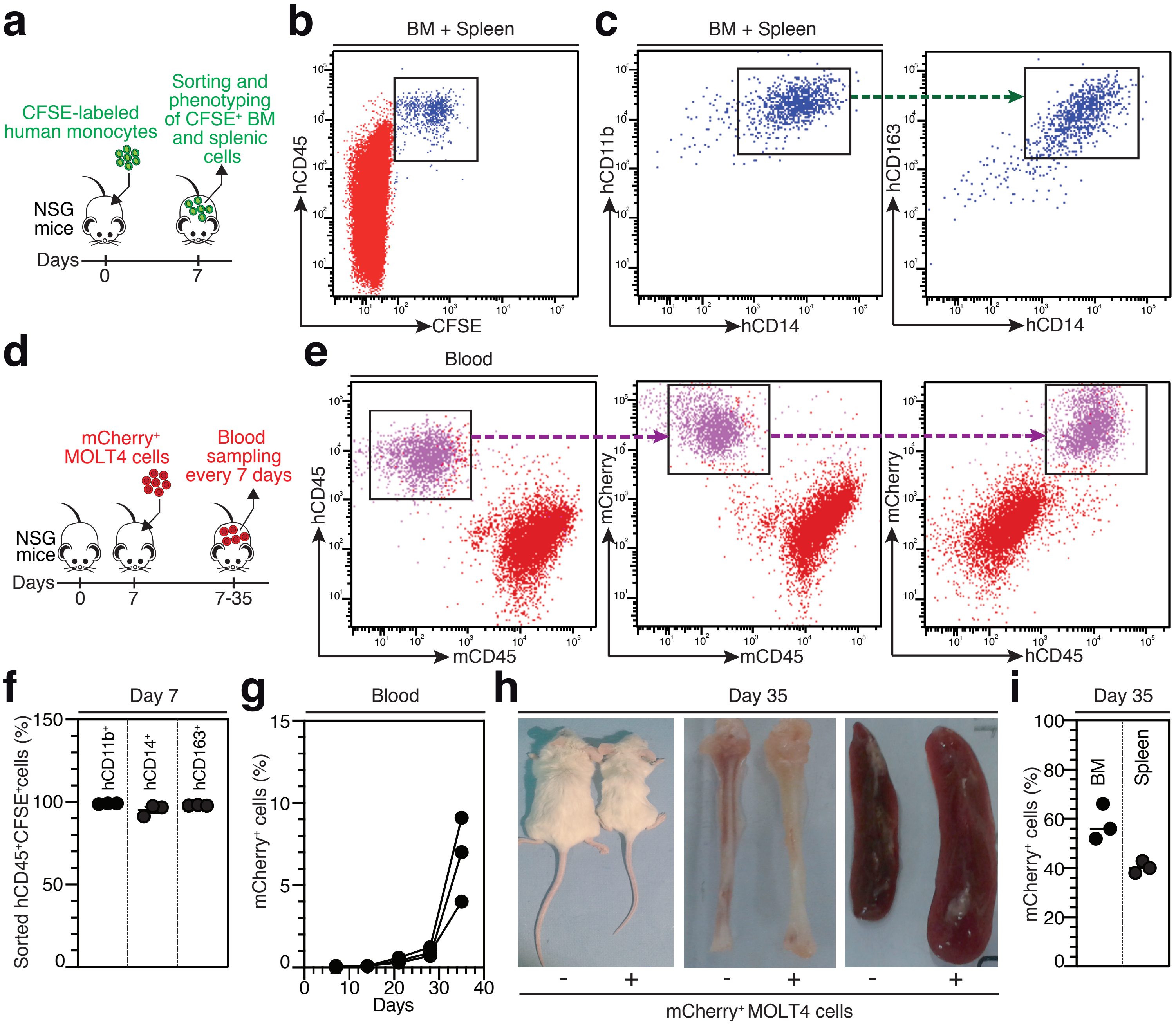


ALLOUCH#EXTENDED4

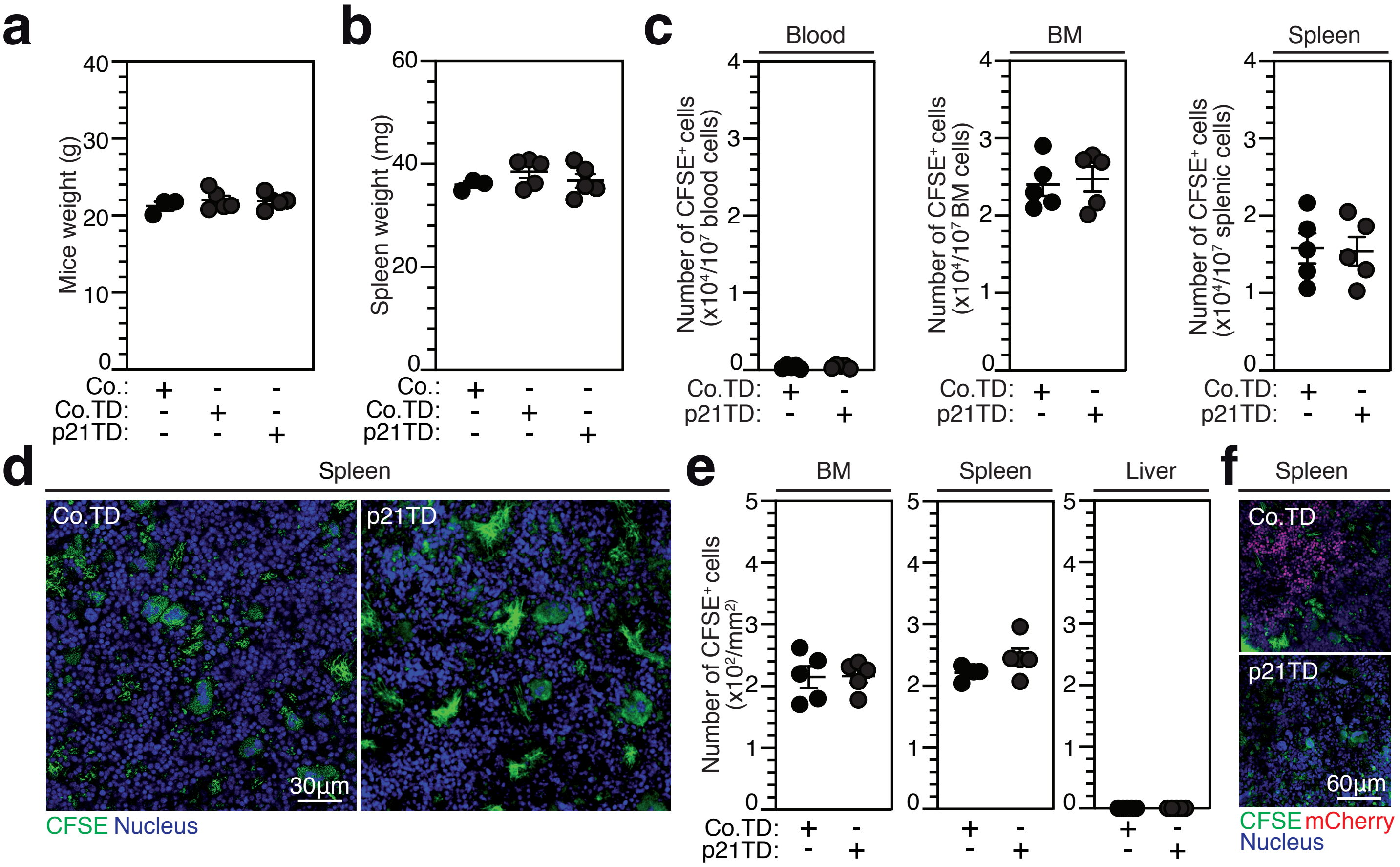


### ALLOUCH#EXTENDED5

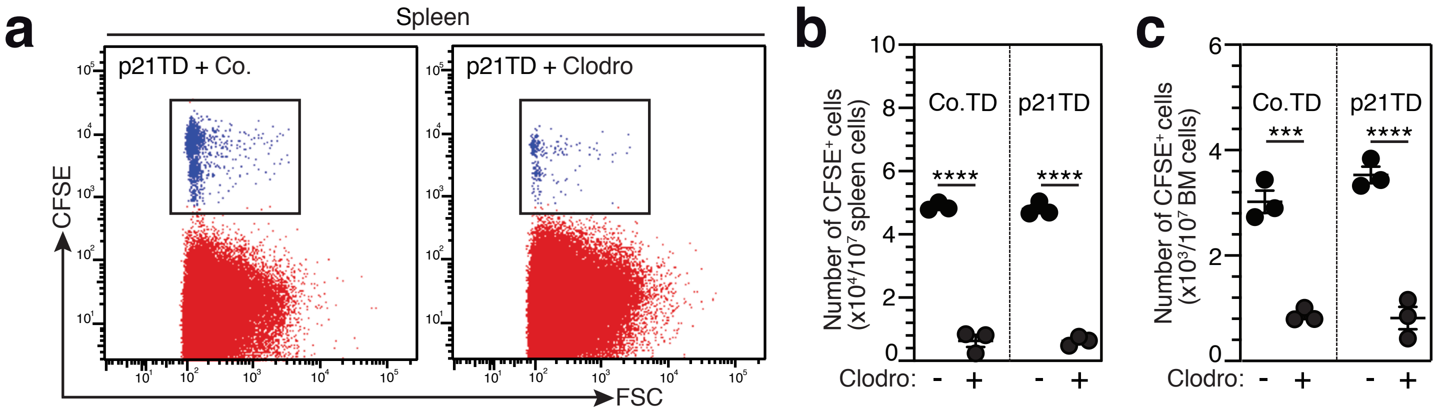


**ALLOUCH#EXTENDED6**

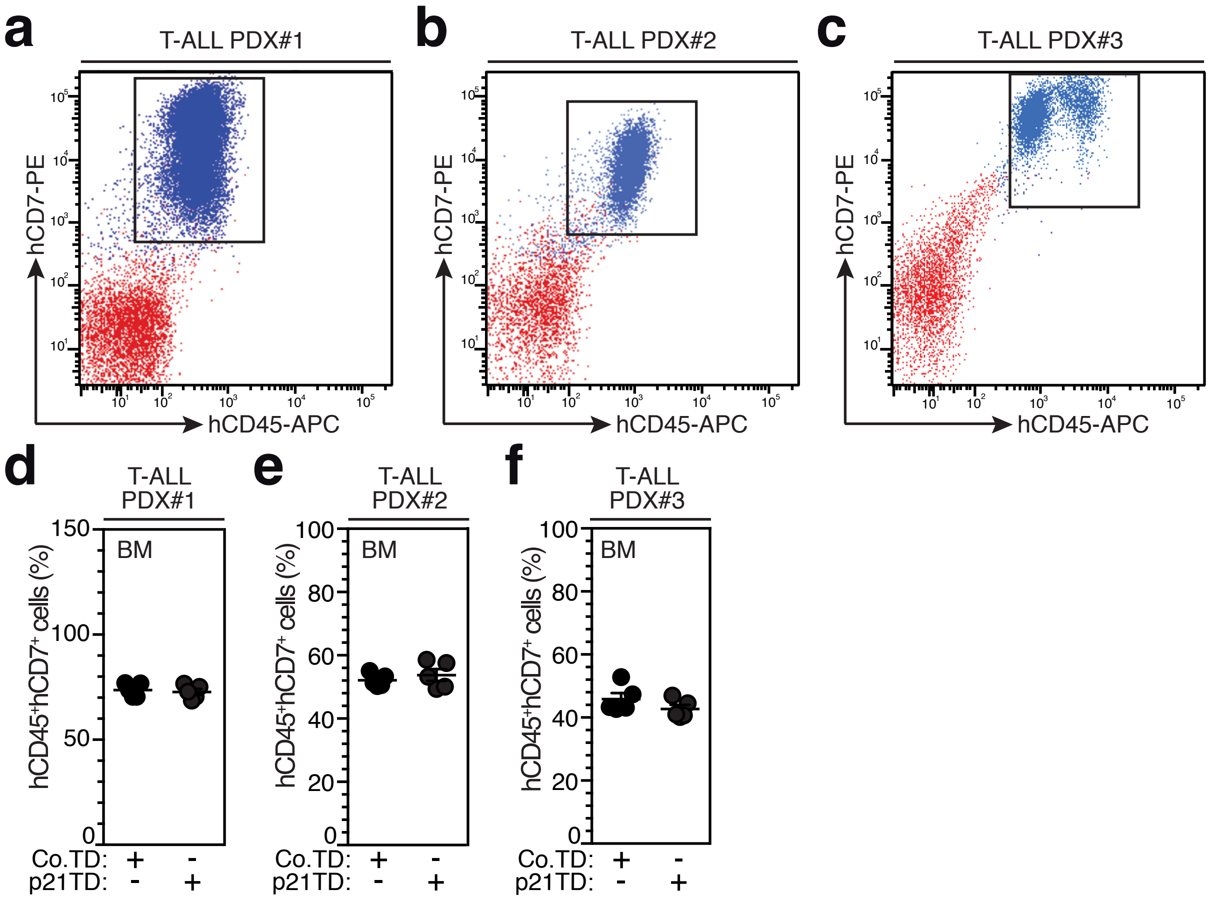


### ALLOUCH#EXTENDED7

**Extended Data Table 1** Characteristics of AML patients of $CD34^+$ cells used in this study

| | Age at sampling time (years) | Sex | WHO diagnosis | WHO diagnosis before AML | Sampled organs | WBC ($10^9$/L) | WBC monocytes ($10^9$/L) | Blasts In BM (%) | Karyotypes | Additional gene mutations | Multi-agent received treatments |
|---|---|---|---|---|---|---|---|---|---|---|---|
| 1 | 73 | Men | AML | CMML | PB | 17.70 | 5 | 79 | 46,XY[20] | *TET2*, *RUNX1 SRSF*2 | Hydroxyurea<br>Hypomethylating agent 5 (HMA)<br>Low cytarabine |
| 2 | 87 | Woman | AML | MDS (RAEB) | BM | 68.20 | 12.96 | 20 | 46, XX[20] | n.d. | Erythropoietin<br>Hypomethylating agent 5 (HMA) |
| 3 | 69 | Men | AML | CMML | PB | 28 | 5.32 | 38 | 46, XY, del(20) (qq11q13) [44]/46, sl, t(8; 21) (q22; q22) [5]/46, sd11, der (22) t(8; 22) (q21-q22; p12) [11] | *ASXL1*, *SRSF2* | Hypomethylating agent 5 (HMA)<br>AML-type chemotherapy |
| 4 | 65 | Men | AML | CMML | PB | 17.9 | 5.7 | 25 | 46, XY [20] | *RUNX1 ASXL1*, *ZRSR2*, *CSF3R* | AML-type chemotherapy<br>Hypomethylating agent 5 (HMA) |
| 5 | 76 | Men | AML | None | BM | 36.40 | 6.55 | 59 | 46, XY[20] | n.d. | Hydroxyurea |
| 6 | 68 | Men | AML | MPN *Polycythemia vera* | PB | 43 | 5.59 | 64 | 46, XY[22] | *JAK2* | Hypomethylating agent 5 (HMA) |

WHO: World health organization; AML: Acute myeloid leukemia; CMML: Chronic myelomonocytic leukemia; MDS: Myelodysplastic syndromes; RAEB: Refractory anemia with excess of blasts (AREB); MPN: Myeloproliferative neoplasms; WBC: Whole blood cells; PB: peripheral blood; BM: Bone marrow; del: deletion; t: translocation; n.d.: not determined

**Extended Data Table 2** List of up-regulated and down-regulated gene expressions in Phago$^{+}$MDMs as compared to Phago$^{-}$MDMs (in Fig. 1j)

| Accession number | Gene Name | Donor-1 Log2FC-1 | Donor-1 Log2FC-2 | Donor-2 Log2FC-1 | Donor-2 Log2FC-2 | Donor-3 Log2FC-1 | Donor-3 Log2FC-2 | Mean Log2FC | Standard deviation | References |
|---|---|---|---|---|---|---|---|---|---|---|
| NM_001243962 | HLA-DQB1 | -0.581085461 | -0.51949771 | -0.543775531 | -0.402907544 | -0.900165827 | -0.901591057 | -0.641503862 | 0.209554892 | 1 |
| NM_001031804 | MAF | -0.748971298 | -0.757355501 | -0.195699637 | -0.154685012 | -0.751059946 | -0.651663441 | -0.543239139 | 0.288053769 | 2 |
| NM_006895 | HNMT | -1.153512746 | -1.312527881 | -0.182960439 | -0.14665632 | -0.728895472 | -0.878054827 | -0.733767947 | 0.485814816 | 3 |
| NM_002003 | FCN1 | -0.441026188 | -0.416885858 | -0.230801825 | -0.318938384 | -0.208387703 | -0.141453228 | -0.292915531 | 0.119938729 | 4,5 |
| NM_001946 | DUSP6 | -0.051145628 | -0.215729214 | -0.270050542 | -0.170089954 | -0.548420144 | -0.465961842 | -0.286899554 | 0.187098811 | 6 |
| NM_001001547 | CD36 | -0.495976027 | -0.52489668 | -0.493696546 | -0.487506267 | -0.550731852 | -0.520427366 | -0.51220579 | 0.024203199 | 3,7 |
| NM_033554 | HLA-DPA1 | -0.968770986 | -0.99228559 | -0.138121342 | 0.005934016 | -1.482841483 | -1.369283679 | -0.824228177 | 0.62288117 | |
| NM_002122 | HLA-DQA1 | -0.852073536 | -0.773154495 | -0.635247778 | -0.765620669 | -0.972414139 | -0.997311877 | -0.832637082 | 0.1371314 | 8 |
| NM_153811 | SLC38A6 | -0.496021681 | -0.470885344 | -0.447700281 | -0.370343873 | -0.541073635 | -0.658669451 | -0.497449044 | 0.09719409 | 3 |
| NM_004244 | CD163 | -1.496105316 | -1.650303955 | -0.580943975 | -0.544766047 | -1.300798552 | -1.113780799 | -1.114449774 | 0.464046325 | 9,10 |
| NM_001150 | ANPEP | -1.07333231 | -1.156206103 | -0.371499456 | -0.59527098 | -0.536939154 | -0.441411901 | -0.695776651 | 0.334583748 | 11,12 |
| NM_005623 | CCL8 | -0.763508565 | -0.788545586 | -0.906444943 | -0.667630277 | -0.564061212 | -0.508529916 | -0.69978675 | 0.148782288 | 13 |
| NM_018092 | NETO2 | -1.340147798 | -1.292177359 | -0.316902202 | -0.332758573 | -1.185540585 | -1.036854232 | -0.917396792 | 0.47069006 | 14,15 |
| NM_004235 | KLF4 | -0.518527327 | -0.430745078 | -0.28226564 | -0.08419777 | -0.137030314 | -0.170882209 | -0.270608056 | 0.173078469 | 16,17 |
| NM_138711 | PPARG | -0.606230518 | -0.627521049 | -0.687632794 | -0.690680448 | -0.388590406 | -0.436133474 | -0.572798115 | 0.129464495 | 18,19 |
| NM_002124 | HLA-DRB1 | -0.631505148 | -0.612412789 | -0.371562307 | -0.344473955 | -0.986503653 | -0.936111262 | -0.647094852 | 0.271172379 | 1,18 |
| NM_019111 | HLA-DRA | -0.394418097 | -0.417530794 | -0.302401288 | -0.32702838 | -0.619917993 | -0.524004868 | -0.43088357 | 0.12100183 | 5 |
| NM_002982 | CCL2 | -4.034458272 | -4.122824587 | -0.330785733 | -0.355001943 | -0.277198599 | -0.285232608 | -1.567583624 | 1.945470015 | 5 |
| NM_005755 | EBI3 | -0.49050625 | -0.628082925 | -0.164509195 | -0.211666767 | -0.881780624 | -0.913493255 | -0.548339836 | 0.320904403 | 20,21 |
| NM_001562 | IL18 | -0.296245809 | -0.343382573 | -0.1780507 | -0.167052357 | -0.45377753 | -0.444585523 | -0.313849082 | 0.124775453 | 22,23 |
| NM_002984 | CCL4 | -1.262445528 | -1.259288658 | -0.583295898 | -0.585440701 | -0.080994996 | -0.074718382 | -0.641030694 | 0.530880867 | 12 |
| NM_002852 | PTX3 | -2.690391917 | -2.724744491 | -1.292646305 | -1.309115891 | -1.453141202 | -1.402132423 | -1.812028705 | 0.696285375 | 24 |
| NM_003596 | TPST1 | -0.774306308 | -0.674300928 | -0.123652374 | -0.092013878 | -0.164570028 | -0.046455513 | -0.312549838 | 0.322832918 | 2 |
| NM_000129 | F13A1 | -1.142032748 | -1.037272543 | -0.038868858 | -0.131280679 | -1.497818827 | -1.312429134 | -0.859950465 | 0.620928529 | 2,25 |
| NM_002731 | PRKACB | -1.011072338 | -1.032191533 | -0.223821059 | -0.384355538 | -0.521610881 | -0.425795189 | -0.599807756 | 0.34066848 | 26,27 |
| NM_001482 | GATM | -0.340103432 | -0.446443832 | -0.601126918 | -0.621274302 | -1.441684205 | -1.437187712 | -0.814636733 | 0.494870001 | 28 |
| NM_000677 | ADORA3 | -1.57763162 | -1.591203995 | -0.356265318 | -0.370834656 | -1.649875119 | -1.92649118 | -1.245383648 | 0.694675261 | 2,27 |
| NM_020362 | PITHD1 | -0.656463418 | -0.608586573 | -0.132963386 | -0.198773236 | -0.464364381 | -0.441486462 | -0.417106243 | 0.212221943 | 9 |
| NM_013308 | GPR171 | -1.062439912 | -0.971339566 | -2.701184225 | -2.470683152 | -0.683938626 | -0.820587156 | -1.45169544 | 0.891018294 | 29 |
| NM_002371 | MAL | -0.843058848 | -0.704240041 | -3.041125348 | -2.943123983 | -0.8856428 | -1.049665515 | -1.577809422 | 1.101488978 | |
| NM_001178126 | IGLL5 | -1.226899044 | -1.011679171 | -2.538817678 | -2.613049755 | -1.501288483 | -1.488248219 | -1.729997058 | 0.680266297 | |
| NM_006159 | NELL2 | -0.675280278 | -0.507305661 | -2.334335332 | -2.488018985 | -0.726561502 | -0.647794551 | -1.229882718 | 0.919193428 | |
| NM_002258 | KLRB1 | -0.554364779 | -0.740102335 | -2.00808683 | -2.136960424 | -1.071601332 | -1.250285814 | -1.293566919 | 0.652033148 | |
| NM_014767 | SPOCK2 | -1.704071577 | -1.857451462 | -2.636960299 | -2.701015041 | -1.668554471 | -1.717989094 | -2.047673657 | 0.485959639 | |
| NM_198196 | CD96 | -0.696125208 | -0.593621022 | -2.231885364 | -2.09452307 | -0.842749897 | -0.825060622 | -1.213994197 | 0.742101926 | |
| ENST00000390237 | IGKC | -1.610846846 | -1.291948829 | -2.789431718 | -2.773346202 | -2.07316802 | -2.035264304 | -2.095667653 | 0.60427258 | |
| NM_001768 | CD8A | -0.662776678 | -0.89732213 | -2.535116732 | -2.411008205 | -0.956368469 | -1.062611052 | -1.420867211 | 0.826418592 | 30 |
| NM_002341 | LTB | -1.613354977 | -1.225439473 | -2.85302617 | -2.94015058 | -1.372357878 | -1.403359095 | -1.901281362 | 0.78133449 | 31 |
| NM_014207 | CD5 | -0.794298307 | -0.722912376 | -2.087569779 | -2.106745623 | -0.671205058 | -0.678643314 | -1.176895743 | 0.714200006 | 32 |
| NM_002104 | GZMK | -0.444578248 | -0.150299334 | -2.282560308 | -2.427657263 | -0.529490154 | -0.621743696 | -1.076054834 | 1.004330829 | |
| NM_014722 | FAM65B | -0.620388577 | -0.219090372 | -2.008572585 | -1.802580879 | -0.268834554 | -0.481008858 | -0.900079304 | 0.794899899 | |
| ENST00000390547 | IGHA1 | -0.374010217 | -0.390645917 | -2.162315619 | -2.118474601 | -1.39912612 | -1.428199573 | -1.312128674 | 0.790347938 | |
| NM_148965 | TNFRSF25 | -1.440527616 | -1.451045121 | -3.083101823 | -3.019569571 | -1.30307197 | -1.350649733 | -1.941327639 | 0.861823909 | |
| NM_006725 | CD6 | -1.19371842 | -0.949347873 | -2.047800435 | -2.040130221 | -0.679723787 | -0.536437995 | -1.241193122 | 0.661554374 | 33 |
| NM_001767 | CD2 | -2.793477034 | -2.657363371 | -4.380364013 | -4.515877283 | -2.078919972 | -2.167563424 | -3.098927516 | 1.081291773 | 34 |
| NM_017933 | PID1 | -1.334882199 | -1.123752236 | -0.182140465 | -0.249230214 | -2.330043267 | -2.701564186 | -1.320268761 | 1.040087832 | 5 |
| ENST00000390551 | IGHG3 | -0.295267099 | -0.367642431 | -0.710481179 | -0.691586762 | -0.341380192 | -0.382887578 | -0.464874207 | 0.185426285 | 35 |
| NM_152866 | MS4A1 | -0.669085636 | -0.593621022 | -0.965302399 | -0.812678376 | -0.749724315 | -0.562576721 | -0.725498078 | 0.150207002 | |
| NM_003121 | SPIB | -0.43060032 | -0.330818667 | -1.079752434 | -0.555633815 | -0.849450814 | -0.777775329 | -0.670671896 | 0.281832671 | |
| NM_006235 | POU2AF1 | -0.846333813 | -1.024642717 | -1.504356826 | -1.609182871 | -1.184297532 | -1.137936986 | -1.217791791 | 0.289168721 | 36 |
| NM_152296 | ATP1A3 | 1.065830065 | 1.153916169 | 1.079159224 | 0.93499379 | 0.380491665 | 0.226792188 | 0.80686385 | 0.399085765 | 37,38 |
| NM_004633 | IL1R2 | 1.6867575 | 1.887059922 | 0.378575927 | 0.203046934 | 0.955640799 | 0.860995048 | 0.995346022 | 0.678376502 | 39,40 |

| | | | | | | | | | | |
|---|---|---|---|---|---|---|---|---|---|---|
| NM_001172 | ARG2 | 1.277183548 | 1.287324072 | 0.312664826 | 0.366667991 | 0.355630197 | 0.512124012 | 0.685265774 | 0.467293172 | 41,42 |
| NM_004615 | TSPAN 7 | 0.677530199 | 0.833303 | 2.512598751 | 2.533954705 | 0.896725707 | 0.912759954 | 1.394478719 | 0.878332776 | 11 |
| NM_002309 | LIF | 0.92385174 | 0.855900108 | 0.134783983 | 0.067470431 | 0.486896914 | 0.616873895 | 0.514296179 | 0.357598844 | 43,44 |
| NM_0010253 66 | VEGFA | 1.052609135 | 1.081569247 | 0.614795091 | 0.587387906 | 1.000890123 | 0.861532163 | 0.866463944 | 0.219196574 | 45,46 |
| NM_000073 | CD3G | 0.683562372 | 0.722470193 | 0.395664755 | 0.563994874 | 0.2211598 | 0.155881262 | 0.457122209 | 0.23803691 | 47,48 |
| NM_000732 | CD3D | 1.939773492 | 2.114086946 | 1.432574455 | 1.507153427 | 1.167837383 | 1.255093409 | 1.569419852 | 0.378549184 | 48,49 |
| NM_000576 | IL1B | 0.120703651 | 0.117081911 | 0.352773716 | 0.294523198 | 1.290071109 | 1.278331349 | 0.575580822 | 0.556798987 | 50,51 |
| NM_0010085 40 | CXCR4 | 1.375972405 | 1.343364601 | 0.88041326 | 0.853668985 | 0.871708008 | 0.900460289 | 1.037597925 | 0.250141408 | 52,53 |
| NM_015714 | G0S2 | 0.981400318 | 0.960626553 | 1.040424186 | 1.01521941 | 2.245997479 | 2.028016295 | 1.37861404 | 0.592114454 | 54 |
| NM_021181 | SLAMF 7 | 0.697913337 | 0.910389113 | 0.064891268 | 0.08679108 | 0.329917011 | 0.244746172 | 0.389107997 | 0.342914175 | 55,56 |
| NM_003486 | SLC7A 5 | 1.727712248 | 1.676075005 | 0.473277137 | 0.510149746 | 0.586725239 | 0.546154357 | 0.920015622 | 0.607028774 | 57 |
| NM_004417 | DUSP1 | 0.835203553 | 0.858259885 | 0.242787469 | 0.223760348 | 0.989660702 | 0.972497069 | 0.687028171 | 0.35674138 | 58,59 |
| NM_175839 | SMOX | 0.46750539 | 0.455103638 | 0.303217854 | 0.349663856 | 0.779522713 | 0.734016178 | 0.514838272 | 0.197975365 | 60 |
| NM_002192 | INHBA | 0.640279166 | 0.582721651 | 0.412412774 | 0.449761872 | 1.775233475 | 1.73246749 | 0.932146071 | 0.642092155 | 61,62 |
| NM_003811 | TNFSF9 | 1.264445984 | 1.08025231 | 0.512080969 | 0.481186666 | 1.092475687 | 1.002212273 | 0.905442315 | 0.328181072 | 63,64 |
| NM_0011008 12 | CXCL1 6 | 1.503012272 | 1.468096755 | 0.212266686 | 0.235775919 | 0.629875745 | 0.648671242 | 0.78294977 | 0.575231637 | 50,65 |
| NM_181054 | HIF1A | 0.948406259 | 0.839172103 | 0.399685798 | 0.395663352 | 1.28177456 | 1.283694902 | 0.858066162 | 0.39828027 | 52,66 |
| NM_000963 | PTGS2 | 0.581174412 | 0.509767842 | 0.104279728 | 0.098204146 | 0.161319789 | 0.107354948 | 0.260350144 | 0.223159691 | 41,67 |
| NM_005375 | MYB | 3.499775743 | 3.389197836 | 4.530634632 | 4.841587902 | 2.152346479 | 2.039161628 | 3.408784037 | 1.164013717 | 68 |
| NM_001661 | ARL4D | 2.435439484 | 2.476020048 | 0.434521353 | 0.538697448 | 1.310952908 | 1.239860346 | 1.405915264 | 0.88739048 | 69 |
| NM_0010029 15 | IGFL2 | 2.709616395 | 2.671378259 | 0.217106968 | 0.284296147 | 1.155564423 | 1.290222678 | 1.388030812 | 1.099678956 | 70 |
| NM_182908 | DHRS2 | 3.416933701 | 3.30810239 | 3.799972307 | 3.714671415 | 1.347452045 | 1.439846691 | 2.837829758 | 1.133725689 | 71 |
| NM_002728 | PRG2 | 4.187722242 | 4.262844823 | 0.825942813 | 0.824745112 | 1.239202917 | 1.358372817 | 2.116471787 | 1.647745928 | 72 |
| NM_012250 | RRAS2 | 2.422034083 | 2.348244032 | 0.73511902 | 0.962415028 | 0.73511902 | 0.962415028 | 1.360891035 | 0.800204723 | 73 |
| NM_016269 | LEF1 | 2.515079235 | 2.518435867 | 1.335993192 | 1.283038146 | 1.329125504 | 1.310715492 | 1.715397906 | 0.621001984 | 74,75 |
| NM_020689 | SLC24 A3 | 2.509869054 | 2.487945778 | 0.24417411 | 0.29596393 | 0.855669424 | 0.686652881 | 1.180045863 | 1.047391562 | 76 |
| NM_016459 | MZB1 | 2.583394848 | 2.596940739 | 0.723107334 | 0.847418516 | 0.723107334 | 0.847418516 | 1.386897881 | 0.933715209 | 77 |
| NM_014220 | TM4SF 1 | 3.684623887 | 3.664781802 | 1.28603437 | 1.322213046 | 1.28603437 | 1.322213046 | 2.094316753 | 1.224284799 | |
| NM_002648 | PIM1 | 2.426849072 | 2.423825868 | 0.429656285 | 0.366686471 | 1.503533027 | 1.466063122 | 1.436102308 | 0.907656914 | 78,79 |
| NM_0011450 33 | AG2 | 3.301542123 | 3.32959608 | 1.297876972 | 1.280865528 | 1.297876972 | 1.280865528 | 1.964770534 | 1.046389336 | |
| NR_026597 | DIRC3 | 2.407209551 | 2.404685824 | 0.272469241 | 0.05800101 | 0.792124093 | 0.832259623 | 1.127791557 | 1.033770692 | |
| NM_0011722 92 | NIPAL4 | 3.697098425 | 3.724208584 | 0.386233055 | 0.357622115 | 2.441272989 | 2.528298357 | 2.189122254 | 1.510887475 | |
| ENST0000038 0464 | PLIN2 | 2.870003562 | 2.72541771 | 0.871565548 | 0.813998109 | 1.432938739 | 1.492002491 | 1.700987693 | 0.895128106 | 80 |
| NM_033120 | NKD2 | 2.411773556 | 2.215007278 | 2.978825795 | 2.738906968 | 1.21440627 | 1.288940654 | 2.141310087 | 0.737945891 | 81 |
| NM_002404 | MFAP4 | 2.736404754 | 2.75575098 | 1.847561408 | 1.806745917 | 0.977980575 | 1.007566721 | 1.855335059 | 0.784591106 | |
| NM_021158 | TRIB3 | 3.216638594 | 3.232921566 | 0.211681968 | 0.189167337 | 0.153481855 | 0.188647295 | 1.198756436 | 1.569470086 | 82 |
| NM_139314 | ANGPT L4 | 3.330031124 | 3.44457626 | 1.140097981 | 0.91885867 | 1.183578945 | 1.19680129 | 1.868990712 | 1.180912556 | 83,84 |
| NM_000597 | IGFBP2 | 1.396431888 | 1.434434584 | 2.865212024 | 2.81262923 | 1.226250159 | 1.166577261 | 1.816922524 | 0.798159749 | 85,86 |
| NM_001853 | COL9A 3 | 1.076008673 | 1.19733003 | 3.400763795 | 3.217354319 | 1.457647379 | 1.231997201 | 1.930183566 | 1.076741237 | |
| NM_0010054 64 | HIST2H 3A | 0.233668323 | 0.164024855 | 2.432292745 | 2.451638157 | 1.347452045 | 1.439846691 | 1.344820469 | 1.004551206 | |
| NM_018667 | SMPD3 | 1.538149607 | 1.624892199 | 3.043850324 | 2.911168449 | 0.859509374 | 1.013238869 | 1.83180147 | 0.935792179 | 87 |
| NM_018492 | PBK | 0.354317436 | 0.235393346 | 2.57989656 | 2.694358302 | 0.529973589 | 0.422286034 | 1.136037545 | 1.167226926 | 88,89 |
| NM_004867 | ITM2A | 2.267706361 | 2.28824222 | 2.570112004 | 2.646682524 | 0.152224439 | 0.197676711 | 1.687107376 | 1.180949555 | 90,91 |
| NR_033916 | CCDC2 6 | 1.467317138 | 1.522604404 | 3.902704634 | 3.934413137 | 1.358916834 | 1.179927602 | 2.227647291 | 1.315025728 | 92 |
| NM_0010043 17 | LIN28B | 1.271591361 | 1.399339878 | 3.158333171 | 3.24915891 | 1.364422982 | 1.546020693 | 1.998144499 | 0.938460457 | 93,94 |
| NM_002612 | PDK4 | 1.884871379 | 1.962521516 | 1.552354138 | 1.426121895 | 2.388744452 | 2.28023544 | 1.915808136 | 0.382493573 | 95,96 |

References for anti- or pro-inflammatory genes: 1, 2, 3, 4, 6, 9, 11, 14, 16, 18, 20, 22, 25, 26, 28, 29, 37, 39, 41, 43, 45, 47, 49, 50, 52, 54, 55, 57, 58, 60, 63, 68, 69, 70, 71, 72, 73, 74, 76, 77, 78, 80, 82, 85, 87, 88, 90, 93 and 95.

References for genes modulated by IFNγ: 1, 2, 5, 7, 8, 10, 12, 13, 15, 17, 18, 19, 21, 23, 24, 27, 30, 31, 32, 33, 34, 35, 36, 38, 40, 42, 44, 46, 48, 51, 53, 56, 59, 62, 64, 65, 66, 67, 75, 79, 84, 86, 89, 91, 92, 94 and 96.

**Extended Data Table 3** Characteristics of T-ALL patients of PDX cells used in this study

| **Patients** | **Age at sampling time (years)** | **Sex** | **WHO diagnosis** | **Karyotypes** | **Additional gene mutations** | **Multi-agent received treatments** |
|---|---|---|---|---|---|---|
| PDX#1 | 16 | Men | T-ALL | 46,XY,t(11;14)(p1?3;q11),add(13)(p?11)[8]/46,XY[19] .nuc ish(ABL1,BCR)x2[200],(TCRADx2)(5'TCRAD sep 3'TCRADx1)[117/200] | IL7R[u]<br>NOTCH1[u] | None |
| PDX#2 | 17 | Men | T-ALL | 46,XY,t(11;14)(p1?3;q11),add(13)(p?11)[8]/46,XY[19] .nuc ish(ABL1,BCR)x2[200],(TCRADx2)(5'TCRAD sep 3'TCRADx1)[117/200] | IL7R[u]<br>NOTCH1[u]<br>PTPRGdel<br>HOXA11del<br>CDKN2Adel<br>CDKN2Bdel<br>TCRDdel<br>NF1gain<br>HLFgain | Daratumomab<br>Venetoclax<br>Dexamethasone<br>Nelarabine<br>Vincristine<br>Cytarabine<br>Etoposide<br>GCSF<br>PL triple |
| PDX#3 | 6 | F | T-ALL | 46,XX,t(11;14)(p13;q11) | SIL-TAL1<br>CDKN2Adel<br>RB1del<br>NOTCH1[u] | None |

WHO: World health organization; T-ALL: T-Cell acute lymphoblastic leukemia
